# Design of high-affinity miniprotein antagonist targeting IL-4Rα for IL-4/IL-13 signal blockade

**DOI:** 10.64898/2026.08.09.743794

**Authors:** Yanghan Yu, Ningning Wang, Lingzhi Xu, Hongyun Wang, Ze Zhang, Bowen Yu

## Abstract

IL-4Ra is a key regulatory receptor for type 2 inflammatory responses, signal transduce from IL-4 and IL-13 through binding with IL-13Ra or the gamma c chain to activate the downstream JAK1-STAT6 pathway. IL-4Ra is currently the most successful “golden target” in the field of allergic disease therapeutics. Its representative monoclonal antibody drug, dupilumab, through the dual blockade mechanism of IL-4/IL-13 has pioneered a new era of precision therapy for type 2 inflammation. In our manuscript, we employed large-scale deep learning-based computational design methods to de novo design mini-protein antagonists specific for both human and mouse IL-4Ra. The binding affinity was improved from 22.1 nM to 569 pM through partial diffusion. The design accuracy and binding specificity were verified through X-ray crystallography and biochemical studies. In vitro IL4/IL13 signal blockade assays revealed that de novo designed monomeric mini-protein antagonist exhibited comparable blockade ability to bivalent dupilumab. In vivo pharmacokinetic half-life studies demonstrated that fusion to an HSA-binding domain extended the half-life of the mini-protein antagonist from 2.7 hours to 60.6 hours. The IL-4Ra mini-protein antagonist had excellent expression levels, solubility and thermal stability. The IL4/IL13 signal blockade ability remained unchanged even after being heating to 95 degrees. In conclusion, through large-scale cluster computing and deep learning-based de novo design, we developed well-performed IL-4Ra mini-protein antagonist, and demonstrates certain potential for drug development.

---

Dear Editor,

IL-4Rα is a key regulatory receptor for type 2 inflammatory responses, signal transduce from IL-4 and IL-13 through binding with IL-13Rα1 or the γc chain to activate the downstream JAK1-STAT6 pathway (**Fig. 1A**). IL-4Rα is currently the most successful “golden target” in the field of allergic disease therapeutics. Its representative monoclonal antibody drug, dupilumab, through the dual blockade mechanism of IL-4/IL-13 has pioneered a new era of precision therapy for type 2 inflammation (Le Floc’h et al., 2020).

**Figure 1.**
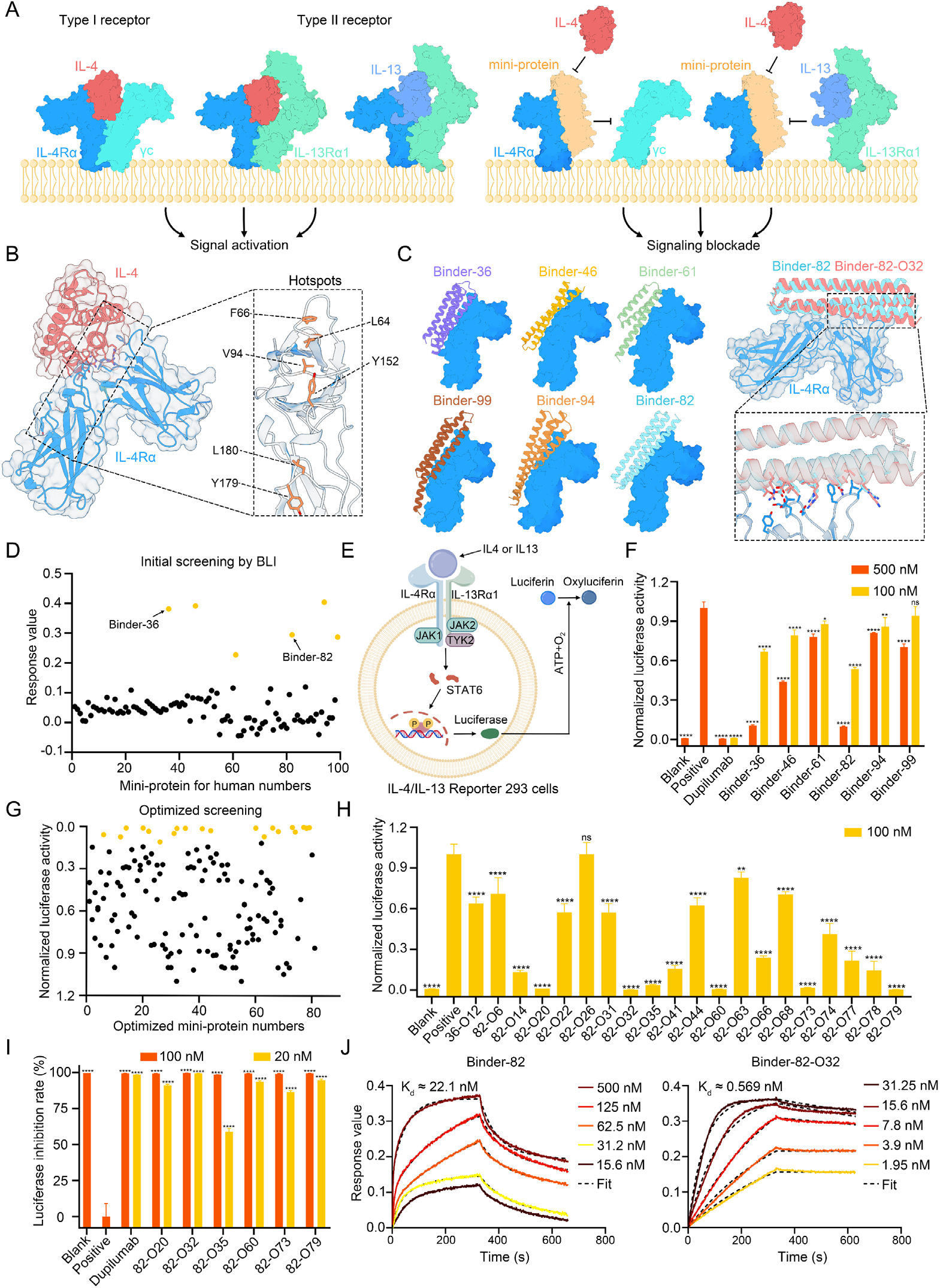
Computational design, screening, and functional characterization of high-affinity IL-4R*α* mini-protein antagonists. **(A)** Schematic diagram illustrating the mechanism by which IL-4Rα mini-protein antagonists block the IL-4/IL-13 signaling pathways. **(B)** Structural interface between IL-4 (red) and IL-4Rα (blue) (PDB: 3BPL). The enlarged view highlights the hotspot residues (orange) selected for binder design. **(C)** Structural snapshots of mini-protein binders obtained from preliminary BLI screening. The parental Binder-82 (cyan) and the partially diffusionoptimized Binder-82-O32 (red) are superimposed, with the inset illustrating interface rearrangements following optimization. **(D)** Preliminary BLI screening results against hIL-4Rα, with the top six candidates highlighted in yellow. **(E)** Schematic diagram of the signaling reporter mechanism in the IL-4/IL-13 Reporter 293 cell line. **(F)** Evaluation of six mini-protein inhibitors for their blockade of IL-4-induced STAT6 signaling pathway activation. **(G)** Functional screening of 150 partially diffusion-optimized mini-protein antagonists (500 nM) for inhibition of 100 pM hIL-4-induced STAT6 signaling using the IL-4/IL-13 Reporter 293 cell line. Candidates with normalized luciferase activity below 0.1 are highlighted in yellow. **(H)** Further assessment of optimized candidates at 100 nM for blockade of 100 pM hIL-4-induced STAT6 signaling. **(I)** Final comparison of six selected candidates at 20 nM for blockade of 100 pM hIL-4-induced STAT6 signaling. **(J)** Binding affinities of Binder-82 and Binder-82-O32 for hIL-4Rα, as determined by BLI. Data in panels **(F), (H)**, and **(I)** are presented as mean ± SD from n = 3 biologically independent samples. ns, not significant, *P < 0.05, **P < 0.01, ***P < 0.001, and ****P < 0.0001.

Despite the remarkable efficacy of IL-4Rα antagonist antibody drugs, chronic diseases require long-term injectable therapy. The relatively high cost limits the accessibility and long-term availability of monoclonal antibody drugs. Compared to traditional antibodies, de novo designed mini-proteins offer significant advantages: purely *in silico* generation, facile *E. coli* expression with high yield, strong thermal stability, high solubility, and flexible engineering properties (Cao et al., 2022). De novo design approaches for mini-proteins have already yielded inhibitors targeting immune components such as TNFR (Glögl et al., 2024), IL-17R (Berger et al., 2024) and C9 (Li et al., 2026). Although these tools are highly adaptable, designing well-performing antagonists against IL-4Rα was still an unmet challenge. Whether mini-proteins can match monoclonal antibody potency while retaining favorable biophysical properties represents a critical test of their druggability.

Blocking IL-4Rα signaling requires disrupting its interaction with IL-4/IL-13(LaPorte et al., 2008). We focused on the cytokine-binding pocket of hIL-4Rα (**Fig. 1B**). In the first round of calculation, we selected four hydrophobic residues L64, F66, V94, and Y152 as hotspots, and generated 16,000 protein scaffolds (50-115 residues) using RFdiffusion (**Fig. 1B**). After ProteinMPNN sequence design and AlphaFold2 complex prediction, screening with pae_interaction < 10 and plddt_binder > 90 yielded a 1.85% success rate (**Fig. S1A**). Analysis revealed that successful binders engaged broader interfaces than cytokines, extending beyond designated hotspots (**Fig. 1B & C**). Notably, successful mini-proteins contained many two-helix binders, empirically poor performers, suggesting insufficient scaffold length. Therefore, in the second round, we added hotspots Y179 and L180 (**Fig. 1B**), generating 16,000 scaffolds with lengths of 60–115 and 115–140 residues. Success rates improved to 2.44% and 5.13% (**Fig. S1B & C**). Mini-proteins with pae_interaction < 6 were further scored through AlphaFold3. Notably, longer mini-protein binders showed better success rates and ipTM values (**Fig. S1D**). Summarizing the above computational results, a total of 100 top ipTM-scoring mini-proteins were selected for experimental screening. We also conducted separate computational screening for mIL-4Rα similarly; results showed substantial differences: successful binders concentrated on the cytokine-binding pocket and were shorter, suggesting limited cross-species activity (**Fig. S3**). For mIL-4Rα, 60 top-ranked candidates were selected for experimental validation.

Following *E. coli* expression and His-tag purification, Bio-Layer Interferometry (BLI) screening identified 6 hIL-4Rα and 3 mIL-4Rα mini-proteins with robust binding (**Fig. 1D & Fig. S3**). Given the greater druggability potential of hIL-4Rα, we focused on the 6 candidates for hIL-4Rα (**Fig. 1C**). We further tested the blocking effects of these 6 candidates in a luciferase-reporter 293 cell line stably expressing IL-4Rα/IL-13Rα1 (**Fig. 1E**). Among these binders, Binder-82 and Binder-36 achieved >85% pathway blockade, yet remained less potent than dupilumab (**Fig. 1F**). We further optimized the affinity of Binder-82 and Binder-36 through partial diffusion.

In partial diffusion optimization, we used Binder-82 and Binder-36 as starting templates, generating 20,000 partial diffusion scaffolds for each binder, followed by similar ProteinMPNN and AlphaFold2 workflows. Models with pae_interaction < 5.4 and plddt_binder > 92 were selected and scored with AlphaFold3. Based on ipTM rankings, 81 and 69 candidates were synthesized for Binder-82 and Binder-36, respectively. After protein expression and purification, candidates were directly screened through the IL-4Rα/IL-13Rα 293 cell line. The first round of screening yielded 20 mini-proteins with complete pathway blockade (**Fig. 1G**). At reduced concentration, six mini-proteins, Binder-82-O20, O32, O35, O60, O73, and O79 could still completely block cell signaling (**Fig. 1H**). Finally, we reduced the concentration of mini-protein binders to

20 nM, finding that only Binder-82-O32 could still completely block the signaling pathway (**Fig. 1I**). Gradient BLI experiments indicated that the affinity of Binder-82-O32 for IL-4Rα improved from 22.1 nM to 569 pM (**Fig. 1J**), demonstrating that partial diffusion significantly enhanced the binding affinity.

To assess design precision, we attempted co-crystallization of IL-4Rα with Binder-82-O32. After incubation and size exclusion chromatography purification, IL-4Rα formed a tight complex with Binder-82-O32 (**Fig. S4**), but well-diffracting crystals were not obtained. We then screened crystals of Binder-82-O32 alone. Due to the strong hydrophilicity of Binder-82-O32, we selected several surface hydrophilic amino acids outside the IL-4Rα binding interface and mutated them to alanine to promote crystal growth (Li et al., 2026) (**Table S2**). We successfully resolved the crystal structure of Binder-82-O32-M2 to 3.5 Å. Structural alignment indicated a Cα r.m.s.d. value of 1.048 Å between the design model and crystal structure, demonstrating high design precision (**Fig. 2A**). Functional assays indicated Binder-82-O32-M2 retained similar inhibitory activity compared to Binder-82-O32 (**Fig. 2B**). To verify the specificity of the binding interface, three interface residues A12, E100 and L111 in Binder-82-O32 were mutated to arginine separately (**Fig. 2A**). Fluorescence-Activated Cell Sorting (FACS) results showed that all three mutations weakened binding to IL-4Rα-positive 293 cells (**Fig. 2C**). Cell signaling blockade experiments also indicated that all three mutations greatly reduced the blocking ability of Binder-82-O32 (**Fig. 2D**).

**Figure 2.**
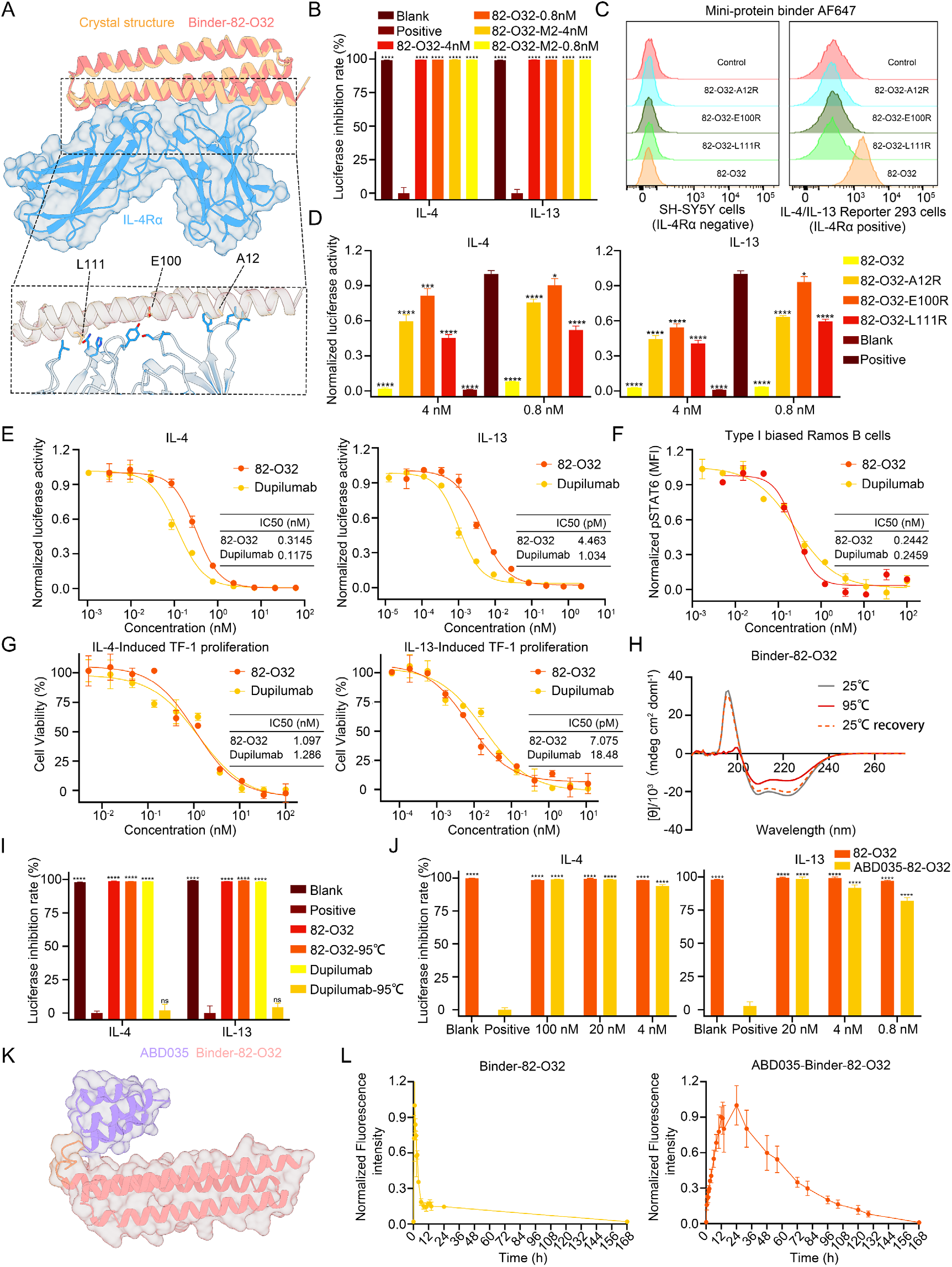
Binding specificity of Binder-82-O32 and comparison of its blocking activity with dupilumab. **(A)** Superposition of the Binder-82-O32-M2 crystal structure (orange) with the predicted Binder-82-O32 (red)/hIL-4Rα (blue) complex model. The enlarged view highlights three key interacting residues in Binder-82-O32. **(B)** Blocking activity of the crystallization construct Binder-82-O32-M2 against 100 pM hIL-4 or 1 nM hIL-13 induced STAT6 signaling in the IL-4/IL-13 Reporter 293 cell line. **(C)** Flow cytometric analysis of the binding specificity of mini-protein antagonists carrying arginine substitutions at different key interaction sites for cell-surface IL-4Rα. Experiments were performed using IL-4/IL-13 Reporter 293 cells with high IL-4Rα expression and SH-SY5Y cells with low IL-4Rα expression. Representative histograms from three independent experiments are shown. **(D)** Blocking activity of Binder-82-O32 variants carrying arginine substitutions at different key interaction sites against 100 pM hIL-4- or 1 nM hIL-13-induced STAT6 signaling. **(E)** Dose-dependent inhibition of IL-4- or IL-13-induced STAT6 signaling in the IL-4/IL-13 Reporter 293 cell line by mini-protein inhibitors and dupilumab. **(F)** Dose-response curves for STAT6 signaling activation in Ramos cells treated with increasing concentrations of dupilumab or Binder-82-O32 in the presence of 1 nM hIL-4, as determined by flow cytometry. **(G)** Dose-response curves for STAT6 signaling activation in TF-1 cells treated with increasing concentrations of dupilumab or Binder-82-O32 in the presence of 5 nM hIL-4 or 5 nM hIL-13. IC_50_ values in panels (e), (f), and (g) were calculated from the corresponding fitted curves. **(H)** CD spectra of Binder-82-O32 under different conditions (gray, 25 °C; red, 95 °C; orange dashed, 25 °C after heat treatment at 95 °C). **(I)** Comparison of the biological activity of Binder-82-O32 before and after heat treatment, with dupilumab used as a reference standard. **(J)** Comparison of the blocking activities of Binder-82-O32 and ABD035-Binder-82-O32 across serially diluted concentrations. **(K)** Structural snapshot of ABD035-Binder-82-O32. **(L)** In vivo serum drug concentrations change of Cy5-labeled Binder-82-O32 and ABD035-Binder-82-O32 in Wistar rats (n = 3). Data in panels **(B), (D), (E), (F), (G) (I), (J)**, and **(L)** are presented as mean ± SD from n = 3 biologically independent samples. ns, not significant, *P < 0.05, **P < 0.01, ***P < 0.001, and ****P < 0.0001.

On this basis, we performed hIL-4Rα gradient signaling pathway blockade assays to directly compare the IC_50_ of Binder-82-O32 with dupilumab. Results showed that the IC_50_ values of Binder-82-O32 and dupilumab for blocking the IL-4-activated signaling were 0.3145 and 0.1175 nM, and for blocking the IL-13-activated signaling pathway were 4.463 and 1.034 pM, respectively, suggesting monomeric IL-4Rα mini-protein inhibitors can achieve comparable effects to bivalent dupilumab (**Fig. 2E**). Given that IL-4/IL-13 Reporter 293 cells predominantly express type II receptor pairs, we subsequently selected Ramos cells, which are biased toward type I receptor expression, for gradient signaling pathway blocking experiments. The results showed that the IC_50_ values of Binder-82-O32 and dupilumab for inhibiting IL-4-induced signaling were highly similar, at 0.2442 and 0.2459 nM, respectively **(Fig. 2F)**. We also examined their inhibitory effects on IL-4- or IL-13-induced proliferation of TF-1 cells, which express both type I and type II receptors. The results also demonstrated that Binder-82-O32 and dupilumab exhibited comparable inhibitory effects **(Fig. 2G)**. Furthermore, we verified the thermal stability of Binder-82-O32. Circular Dichroism (CD) experiments demonstrated that Binder-82-O32 fully recovered its secondary structure after heating to 95°C and returning to room temperature (**Fig. 2H**). Blockade assay results showed that Binder-82-O32 completely retained its inhibitory activity, while dupilumab’s signaling pathway inhibitory activity was completely lost (**Fig. 2I**). Additionally, Binder-82-O32 exhibited high expression levels, reaching ∼60.5 mg/L without optimization (**Fig. S5**).

As a monomeric small mini-protein inhibitor, Binder-82-O32 has limited in vivo half-life which hinders its druggability. We attempted to extend the in vivo half-life of Binder-82-O32 by fusing the HSA-binding domain ABD035(Nilvebrant and Hober, 2013) at the N-terminus. Cell signaling blockade experiments indicated that ABD035 fusion had limited impact on the inhibitory activity of Binder-82-O32 (**Fig. 2J**). Rat in vivo half-life experiments showed that ABD035 fusion successfully extended its half-life from 2.706 to 60.6 hours (**Fig. 2I**).

In conclusion, through large-scale cluster computing and deep learning-based de novo design, we developed well-performed IL-4Rα mini-protein antagonist. Although dupilumab exhibits a stronger binding affinity (<100 pM) (**Fig. S2**), the broader blocking interface enables mini-protein antagonist achieve comparable levels of signaling pathway inhibition to dupilumab. Future engineering to dimerize the HSA-binding domain, forming antibody-like homodimers to enhance blockade while retaining half-life extension, represents a valuable direction. Additionally, validation of the biological effects of mini-protein inhibitors in humanized mouse disease models and immunogenicity evaluation are also necessary.

## Author contributions

Bowen Yu and Yanghan Yu designed the research. Bowen Yu made the designs. Yanghan Yu performed most of the experiments with the help of Ningning Wang, Lingzhi Xu, Hongyun Wang and Zhang Ze. All authors analyzed data. Bowen Yu supervised the research. Bowen Yu and Yanghan Yu wrote the manuscript with the input from the other authors. All authors revised the manuscript.

## Acknowledgements

This work was supported by grants from the National Natural Science Foundation of China (32501303 to Bowen Yu, 82303251 to Ningning Wang, 81701590 to Lingzhi Xu), the Shandong Provincial Natural Science Foundation, China (ZR2022QC209 to Bowen Yu, ZR2023QH202 to Ningning Wang, ZR2024MH338 to Lingzhi Xu). This work was also supported by the Shandong Provincial Science and Technology Support Plan for Youth Innovation in Universities (10438202502 to Bowen Yu). We would like to thank the Protein Characterization and Crystallography Facility of Westlake University for help in sample analysis; the Mass Spectrometry & Metabolomics Core Facility of Westlake University for sample analysis; and the Westlake University HPC Center for computation assistance.

## Data availability

Coordinate and structure file has been deposited to the Protein Data Bank with accession codes 24WD [https://doi.org/10.2210/pdb24WD/pdb]. The final sequences of the designed inhibitors optimized by partial diffusion are provided in **Table S2**. Full raw data including all tested models will be available on zenodo.org.

