## supplementary data for "Design of high-affinity miniprotein antagonist targeting IL-4Rα for IL-4/IL-13 signal blockade"

### Methods

#### Ethics statement

All animal experiments were conducted in accordance with the guidelines and regulations of the Laboratory Animal Management Committee of Shandong Province, China, and were approved by the Experimental Animal Ethics Committee of Shandong Second Medical University (No. 2025SDL799). Wistar rats were maintained under specific pathogen-free (SPF) conditions with a 12-h light/dark cycle and were provided with standard laboratory chow and water. The animal facility was maintained at a temperature of 20–24 °C and a relative humidity of 50%–60%.

#### Computational binder design

The binder design workflow followed previously published protocols (Watson et al., 2023; Glögl et al., 2024; Li et al., 2026). RFdiffusion, ProteinMPNN, AlphaFold2, and AlphaFold3 were implemented on a local computing cluster without modification of the original source code. The crystal structure of IL-4Rα (PDB: 3BPL) was selected as the initial target. For each RFdiffusion-generated backbone, three sequences were designed using ProteinMPNN. Partial diffusion was performed with diffuser.partial_T values of 6, 8, 10, 11, 12, 13, 14, 15, 16, 17, 18, 19, 20, 21, 23, and 25, generating 1,250 scaffolds for each T value. These scaffolds were subsequently subjected to the same ProteinMPNN sequence design and AlphaFold2 structure prediction pipeline. The final sequences of both the initially screened and partial diffusion-optimized mini-protein binders are provided in **Table S2**.

#### Protein expression and purification

Mini-protein binder coding sequences were codon-optimized for expression in *E. coli*. The corresponding genes were synthesized and cloned into the pET-28a vector between the NcoI and XhoI restriction sites. The hIL-4Rα ectodomain (M26-E227) was codon-optimized for expression in *H. sapiens*. The corresponding sequence was fused with an

N-terminus mouse Ig kappa secretion signal and a C-terminal $(His)_6$, and inserted between the XbaI and BamHI sites of the pcDNA3.4 vector. All genes were synthesized and constructed by Universe Gene Technology or GENCEFE Biotech.

Plasmids encoding mini-protein binders were transformed and expressed in *E. coli* BL21(DE3). Briefly, the transformed single colony was inoculated in 100 ml LB medium containing 50 μg/mL kanamycin, and cultured at 37 °C with shaking at 220 rpm. When the cultures reached an OD600 of 0.6–0.8, IPTG was added to a final concentration of 0.3 mM to induce protein expression for 12–16 h at 24 °C. After expression, E. coli was collected by centrifugation at 10,000 × g. After removing the supernatant, E. coli was resuspended in lysis buffer (PBS), and lysed by ultrasonic disruption. The insoluble impurities were removed from the lysate by centrifugation at 12 000 × g for 20 min. The supernatant was loaded into a pre-balanced Ni-NTA affinity chromatography column for purification; non-target proteins were washed using PBS solutions containing 10, 20, and 30 mM imidazole, respectively. Mini-protein binders were eluted with PBS plus 300 mM imidazole, pH 7.4. All candidate proteins were successfully expressed, except for one candidate from the initial hIL-4Rα screening that failed to express properly.

For hIL-4Rα expression, plasmids were sterile-filtered and transiently expressed using a mammalian expression system. After 5 days of expression, supernatants were harvested. $(His)_6$-tagged hIL-4Rα was purified using Ni Smart beads (Smart-Lifesciences), following the manufacturer's protocols.

For hIL-4Rα mini-protein complex formation, hIL-4Rα was mixed with an excess Binder-82-O32 or Binder-82-O32-M2, incubated at room temperature for 30 min, and subjected to SEC on a Superdex 75 Increase 10/300 GL column (GE Healthcare) using an FPLC system (Union-biotech, UEV25D). Protein concentrations were determined using a multimode microplate reader by measuring the absorbance at 280 nm. Circular dichroism (CD) characterization was conducted following a previously reported protocol (Yu et al., 2025; Li et al., 2026).

**BLI**

BLI experiments were carried out using an OCTET RED96E system (ForteBio), and the results were processed with integrated software (Fortebio Data Analysis 12.0.1.2). All BLI experiments were performed using a standard protocol that included the following steps: baseline, loading, a second baseline, association, and dissociation. All the baseline steps were set to 60 s. The Fc-tagged IL-4RA protein (Novoprotein, CS38; ACRO Biosystems, ILR-M52H1) was used as ligands, diluted to a final concentration of 5 μg/mL before immobilization onto Protein A biosensors (ForteBio) for 600 s. The association and dissociation phases lasted 330 s and 300 s, respectively. For initial screening, mini-protein binders were diluted to 1 μM (**Fig. S2**, **S3A & S3D**). For kinetic binding analyses, mini-protein binders were serially diluted to concentrations ranging from 500 to 1.95 nM (**Fig. 1J & S2**). PBST (PBS containing 0.05% Tween-20, pH 7.4) was used for sensor equilibration, baseline stabilization, and sample dilution. The shaking speed was set to 600 rpm during ligand loading and 1,000 rpm for all other steps. All experiments were performed at 25 °C. The data shown in **Fig. 1J** and **Fig. S2**, which include dupilumab, were fitted globally using a 1:1 fitting model, while **Fig. S2** and **S3A** (excluding dupilumab) were fitted locally using a 1:1 fitting model. Fitting model for the Binder-82 model shown in Figure 1J resulted in $R_{max}$ of 0.3766, $R^2$ of 0.9964, $X^2$ of 0.2161, and $K_d$ of 22.18 nM. Fitting model for the Binder-82-O32 model shown in **Fig. 1J** yielded $R_{max}$ of 0.6492, $R^2$ of 0.9957, $X^2$ of 0.2317 and $K_d$ of 0.569 nM. Fitting model for dupilumab in **Fig. S2** yielded $R_{max}$ of 0.4824, $R^2$ of 0.9967, $X^2$ of 10.4832 and $K_d$ of 7.191 pM. In order to subtract the background, each independent experiment included a reference well with no mini-protein inhibitor added in the association step.

**Cell culture**

IL-4/IL-13 Reporter 293 cells (Genomeditech, GM-C01511) were cultured in DMEM (Gibco) supplemented with 10% fetal bovine serum (FBS; Gibco), 1% penicillin–

streptomycin (10,000 U $mL^{-1}$; Gibco), 4 μg $mL^{-1}$ (MCE, HY-K1054), and 0.75 μg $mL^{-1}$ puromycin (Sangon Biotech, E607054-0001). SH-SY5Y and Ramos cells were maintained in RPMI 1640 medium (Gibco) supplemented with 10% FBS. TF-1 cells were cultured in RPMI 1640 medium containing 10% FBS and 0.2 ng $mL^{-1}$ GM-CSF (Sino Biological, 10015-HNAN). All cells were maintained at 37 °C in a humidified incubator with 5% $CO_2$, were used for no longer than 2 months after thawing, and were routinely tested for mycoplasma contamination.

**FACS**

To evaluate the binding activity of mini-proteins and their inhibitory effects on the IL-4 signaling pathway, the following two experiments were conducted:

Binding activity assay: IL-4/IL-13 Reporter 293 and SH-SY5Y cells ($1 \times 10^5$ cells/well) were seeded into 24-well plates and cultured overnight. Cells were then incubated with mini-protein binders at 4°C for 30 min; control groups received an equal volume of PBS. Following incubation, cells were washed three times with PBS to remove unbound proteins. Subsequently, cells were stained with rabbit anti-6×His monoclonal antibody (Proteintech, CL647-66005) at 4°C for 30 min. After three additional washes, cells were resuspended in PBS and subjected to flow cytometry analysis. The binding intensity was quantified as mean fluorescence intensity (MFI).

Signaling pathway inhibition assay: Ramos cells were seeded into 24-well plates ($2 \times 10^5$ cells/well) and serum-starved overnight in RPMI 1640 medium supplemented with 2% FBS. The following day, 3-fold serially diluted dupilumab and mini-proteins were mixed with hIL-4 (1 nM) and added to the corresponding wells. After gentle pipetting to mix, cells were incubated at 37°C in a 5% $CO_2$ incubator for 15 min. Subsequently, 300 μL of freshly prepared 4% paraformaldehyde (PFA) (Biosharp, BL539A) was added to each well to achieve a final concentration of 1.5%; after vortex mixing, cells were fixed at room temperature for 10 min. Cells were then centrifuged at 4°C, 375 × g for 5 min, and the supernatant was discarded. After thorough resuspension of the cell pellet in the residual supernatant, 1 mL of ice-cold methanol was added to each tube. Cells were incubated at 4°C for 30 min and were washed with 3 mL of FACS buffer, centrifuged at 4°C, 375 × g for 5 min, the supernatant was discarded; An appropriate volume of FACS buffer was added to resuspend the cells, followed by staining with anti-Hu/Mo Phospho-STAT6 antibody (Invitrogen, 17-9013-42). Cells were incubated at room temperature in the dark for 30 min and then subjected to flow cytometry analysis. The phosphorylation level was quantified as median fluorescence intensity (MFI). Normalized pSTAT6 (MFI) = (sample MFI − blank control MFI) / (positive control MFI − blank control MFI).

FACS was performed using a BD FACSAria III instrument (BD Biosciences); Data were analyzed using FlowJo software (version X10.0.7r2).

**Crystallization and structural determination**

Initial crystal screening was conducted using kits purchased from Hampton Research and Rigaku, which included Crystal Screen, PEGRx, and Wizard Classic 1-4. Initial screenings, along with subsequent optimizations, were carried out using a two-position deck mosquito LCP (SPT Labtech) in 96-well crystallization plates using the sitting-drop method. To improve the crystal quality, 3–4 surface alanine mutations were introduced into mini-protein inhibitors. Sequences are shown in **Table S2**. The $(His)_6$ tag and designed sequences were directly fused. Mini-proteins used for crystal screening were further purified by SEC and ultrafiltered to a concentration of ~75 mg/mL. For hIL-4Rα mini-protein complex crystal screening, the complex fraction was concentrated to ~15 mg/mL. The crystallization conditions for Binder-82-O32-M2 were 6% v/v 2-Propanol, 0.1 M Sodium acetate trihydrate pH 4.5, 26% v/v Polyethylene glycol monomethyl ether 550. The crystals were subjected to diffraction and indexing using an in-house X-ray diffraction system (Rigaku) utilizing Cu Kα radiation with a wavelength of 1.54 Å. The structure was solved using the molecular replacement method, employing the designed models as search templates. Molecular replacement and structure refinement were executed using the PHENIX suite(Adams et al., 2010) and COOT(Emsley and Cowtan, 2004). Visualization of the structures was performed using ChimeraX(Goddard et al., 2018) and PyMOL (The PyMOL Molecular Graphics

System, 2002). The collection and refinement statistics are comprehensively presented in **Table S1**.

**IL4/IL13 cell signal inhibition assay**

IL-4/IL-13 Reporter 293 cells were seeded at a density of $1.5 \times 10^4$ cells per well in 96-well plates 20–24 hours prior to stimulation and cultured overnight at 37°C with 5% $CO_2$. After removal of the supernatant, serial dilutions of mini-protein inhibitors, recombinant human IL-4 (100 pM; Novoprotein, CX03) or IL-13 (1 nM; Novoprotein, CC89), and Dupilumab (Sanofi) were prepared in DMEM containing 1% FBS (Gibco) and added to the corresponding wells. The experimental groups were set up as follows: experimental wells received mini-protein inhibitors plus cytokine; dupilumab control wells received serially diluted dupilumab plus cytokine; positive control wells received cytokine alone; and blank control wells received DMEM with 1% FBS only. Following incubation at 37°C for 7 hours, 100 μL of Bio-Lumi™ II Firefly Luciferase Reporter Gene Assay Kit (Beyotime, RG043M) was added to each well. After equilibration at room temperature for 5 minutes, the plates were transferred to white opaque detection plates and chemiluminescence was measured using a multimode microplate reader (Tecan Spark).

Candidate proteins identified by BLI screening were initially evaluated for inhibition of IL-4 signaling at concentrations of 500 and 100 nM. For IL-4Rα-targeting optimized inhibitors, a stepwise concentration reduction strategy (500, 100, and 20 nM) was applied during functional screening, leading to the identification of the lead candidate, Binder-82-O32. Subsequent functional characterization was performed using concentration gradients for selected variants, including the crystallographic construct Binder-82-O32-M2 (100 and 20 nM; IL-4), three arginine-substituted mutants (4 and 0.8 nM; IL-4 and IL-13), and the HSA-binding domain-fused construct ABD035-Binder-82-O32 (IL-4: 100, 20, and 4 nM; IL-13: 20, 4, and 0.8 nM).

Dupilumab and Binder-82-O32 were heated at 95°C for 5 minutes, cooled to room temperature, and subsequently tested for IL-4 and IL-13 signaling inhibition at 20 nM to compare activities before and after the heat treatment.

All gradient experiments employed serial dilutions prepared using equal-ratio dilution method. Blank wells were not treated with IL-4 or IL-13. Luciferase activity was normalized and inhibition rates were calculated using the following formulas: Normalized luciferase activity = (sample $_{luminance}$ – blank $_{Luminance}$) / (positive $_{Luminance}$ – blank $_{Luminance}$); luciferase inhibition rate = [1 – (sample $_{luminance}$ – blank $_{Luminance}$) / (positive $_{Luminance}$ – blank $_{Luminance}$)] × 100.

**TF-1 Cell Proliferation Assay**

TF-1 cells in the logarithmic growth phase were washed twice with sterile PBS to remove residual GM-CSF, then resuspended in RPMI 1640 medium supplemented with 2% FBS and starved overnight. Prior to the experiment, cells were resuspended in RPMI 1640 medium containing 10% FBS, and the cell density was adjusted to $4 \times 10^5$ cells/mL for subsequent use. In a 96-well flat-bottom culture plate, antibodies were serially diluted 3-fold in assay medium, with 10 μL added per well. Subsequently, 50 μL of medium containing recombinant human IL-4 (final concentration 80 ng/mL), IL-13 (final concentration 80 ng/mL), or IL-4 plus IL-13 (each at a final concentration of 80 ng/mL) was added to the respective wells, bringing the total volume per well to 60 μL. The plate was pre-incubated in a 37°C, 5% $CO_2$ incubator for 30 min. TF-1 cell suspension (50 μL, $2 \times 10^4$ cells/well) was then added to each well, resulting in a final volume of 110 μL per well. The plate was continuously cultured in a 37°C, 5% $CO_2$ incubator for 72 h. At the end of the culture period, 10 μL of enhanced CCK-8 reagent (Beyotime, C0042) was added to each well, gently mixed, and the plate was further incubated in a 37°C, 5% $CO_2$ incubator for 1–4 h (2–4 h recommended for suspension cells). The absorbance of each well at 450 nm ($OD_{450}$) was measured using a multimode microplate reader (Tecan Spark). TF-1 cell viability was calculated using the following formula:Cell viability = (sample $OD_{450}$ − blank control $OD_{450}$) / (positive control $OD_{450}$

− blank control $OD_{450}$)

**Cy5 covalent labeling**

A mini-protein containing a cysteine mutation was treated with TCEP (Beyotime, ST046) and then labeled with sulfo-Cy5 maleimide (DuoFluor,2242791-82-6). Briefly, the procedure was as follows: The purified mini-protein containing the cysteine mutation (Binder-82-O32-43C, ABD035-82-O32-43C) was mixed with TCEP at a 10:1 molar ratio and reacted overnight at 4°C. After removing TCEP via size exclusion chromatography (SEC), the mini-protein-Cy5 conjugate was collected and excess Cy5 dye was added; the mixture was then incubated at room temperature on a rotary mixer for 2 hours. Upon completion of labeling, unbound Cy5 dye was removed via size exclusion chromatography on a Superdex 75 Increase 10/300 GL column (GE Healthcare) using an FPLC system (Union-biotech, UEV25D). The degree of labeling was further assessed by SDS-PAGE as described below.

**SDS-PAGE**

To detect the degree of labeling of Cy5 on the mini-protein binder, a final concentration of 2 mM CuSO4 was added to the Binder-82-O32-43C or Binder-82-O32-43C-HSA samples, while an equal amount of PBS was added to the control group. The samples were then oxidized at room temperature for 30 min. Subsequently, 10 μg of the protein sample was mixed with 5×loading buffer, heated at 95°C for 5 min, and loaded onto the corresponding lane for electrophoresis.

**Pharmacokinetic Analysis in Rats**

Female SPF-grade Wistar rats (6–8 weeks old, n = 3 per group) received a single subcutaneous injection of Cy5-labeled mini-protein at a dose of 3 mg/kg. Blood samples were serially collected via the retro-orbital venous plexus or tail vein at 0, 0.08, 0.16, 0.25, 0.50, 0.75, 1.00, 2.00, 3.00, 4.00, 6.00, 8.00, 12.00, 24.00 h post-injection. After centrifugation, serum samples were collected, and the fluorescence signal intensity was measured using a multimode microplate reader (Tecan Spark). Plasma concentration-time profiles were constructed, and pharmacokinetic parameters were calculated to characterize the in vivo pharmacokinetic properties of the compound. Normalized fluorescence intensity is calculated as $F_\lambda/F_{max}$, where $F_\lambda$ is the fluorescence intensity at wavelength λ and $F_{max}$ is the maximum fluorescence intensity at the emission peak.

### Statistics

Data in figures were presented as mean ± SD. Statistical analyses were performed using one-way analysis of variance for multiple-group comparisons. The gradient IL-4/IL-13 cell signal inhibition assay used a standard four-parameter dose–response inhibition function to calculate IC50 (50% inhibition) values. For in vivo drug concentration analysis, pharmacokinetic parameters were calculated by fitting the concentration-time data to a one-phase decay model. The terminal half-life ($t_{1/2}$) was determined from the decay.ns, not significant, *$P < 0.05$, **$P < 0.01$, ***$P < 0.001$, and ****$P < 0.0001$. Analysis was carried out using GraphPad Prism 8.4 for Windows (GraphPad Software, www.graphpad.com).

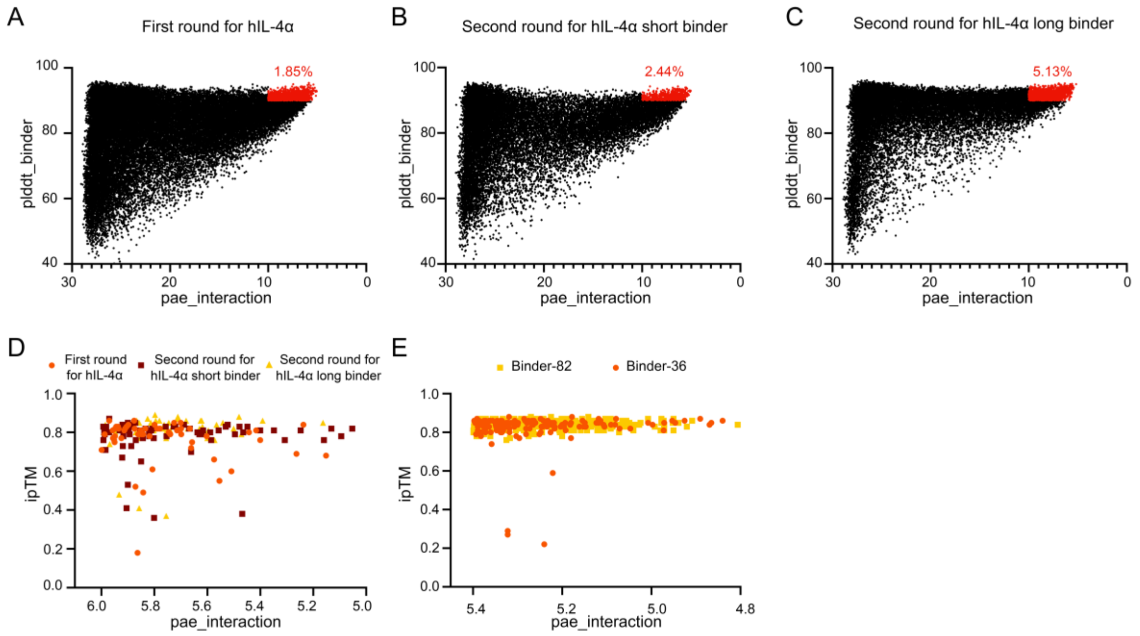

**Supplementary Fig. 1. Calculation scores for mini-protein binding design to hIL-4Rα.** AlphaFold2-predicted scores for the first **(A)**, second **(B)**, and third **(C)** rounds of screening. Scores meeting the threshold criteria of pae_interaction < 10 and plddt_binder > 90 are highlighted in red. **(D)** AlphaFold3 scoring prediction for mini-protein binders with pae_interaction < 6 across all three rounds. **(E)** AlphaFold3 scoring prediction for mini-protein binders derived from Binder-36 and Binder-82 after partial diffusion optimization with pae_interaction < 5.4.

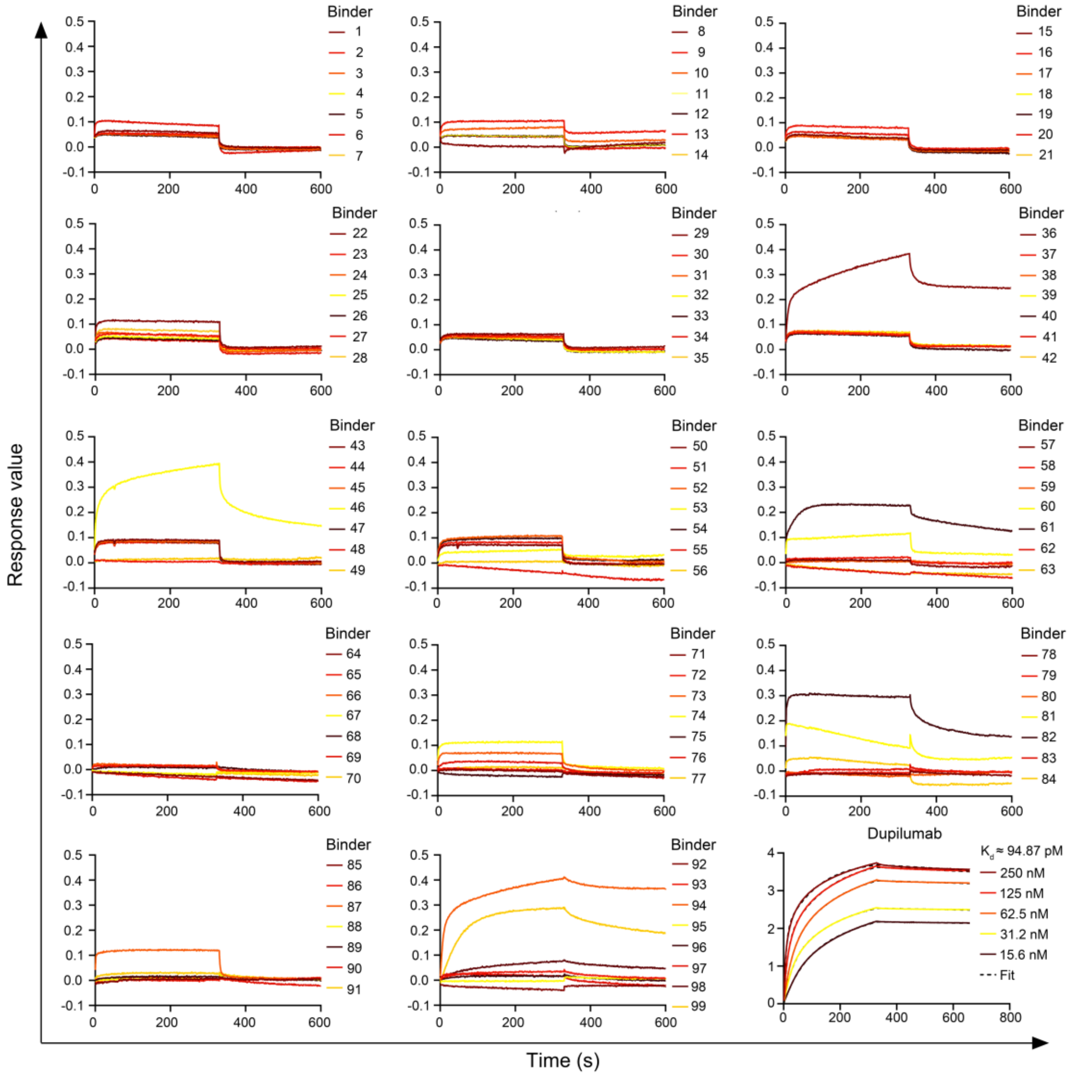

**Supplementary Fig. 2. BLI analysis of mini-protein antagonists targeting hIL-4Rα.** BLI data of the binding between mini-protein inhibitors and hIL-4Rα from the initial screening. The bottom right panel depicts the binding kinetics between dupilumab and hIL-4Rα.

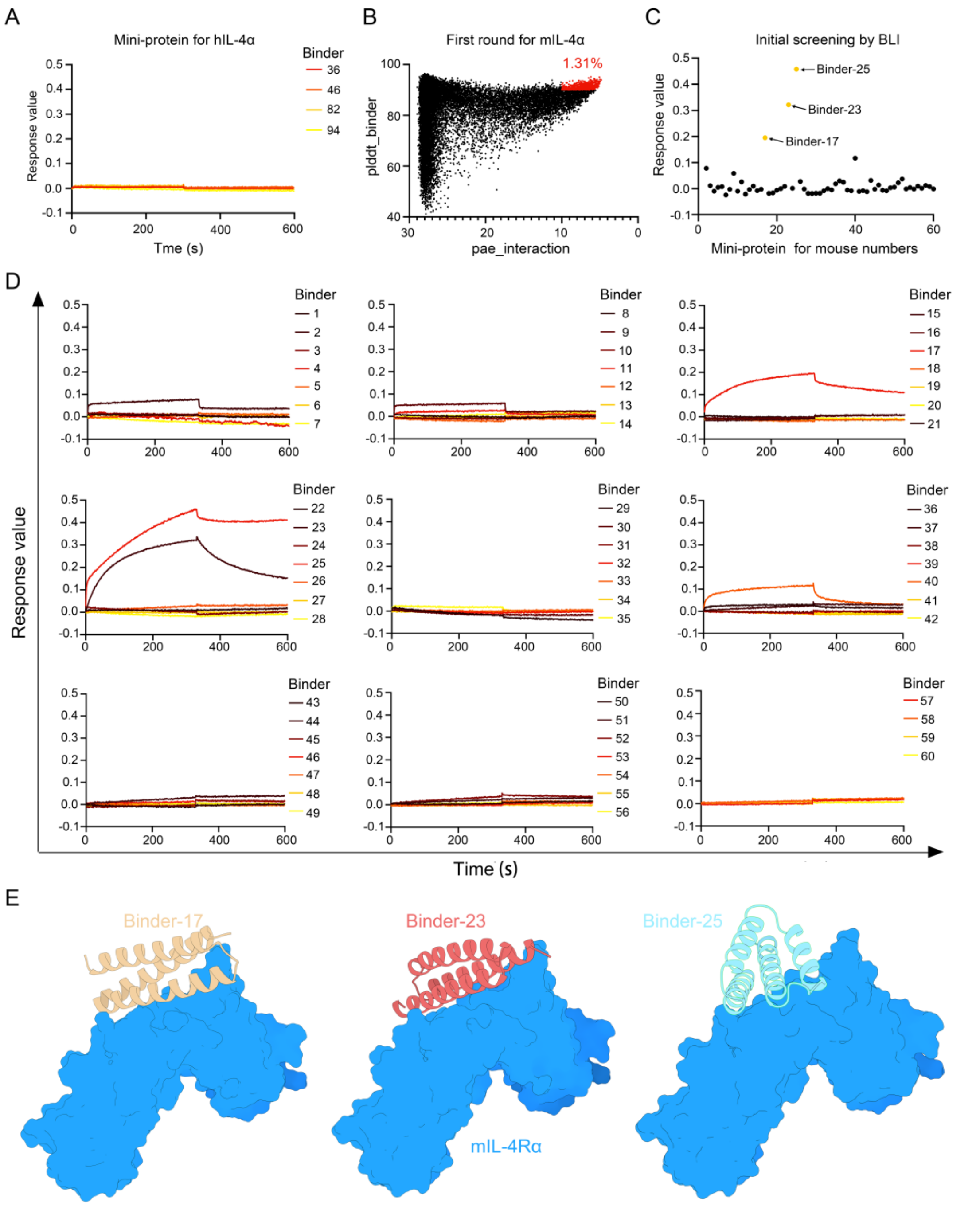

**Supplementary Fig. 3. Design and screening data for mini-protein inhibitors targeting mIL-4Rα**. **(A)** BLI data demonstrated that the four mini-proteins identified from initial screening against hIL-4Rα did not interact with mIL-4Rα. **(B)** AlphaFold2-predicted scores for mini-protein inhibitors targeting mIL-4Rα. Scores meeting the threshold criteria of pae_interaction < 10 and plddt_binder > 90 are highlighted in red. **(C)** Statistical graph of response value in **D**. **(D)** BLI data of the binding interactions between mIL-4Rα and 60 mini-protein inhibitors from the initial screening. **(E)** Structural snapshots of Binder-17, Binder-23, and Binder-25 obtained from preliminary BLI screening.

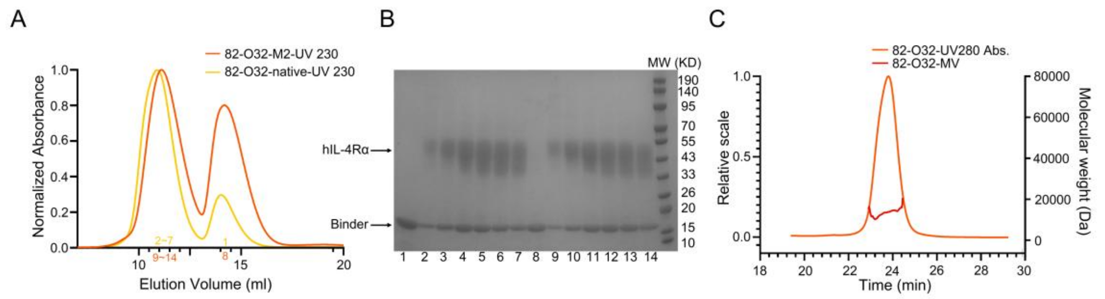

**Supplementary Fig. 4. Complex formation between Binder-82-O32 and hIL-4Rα on SEC. (A)** Size-exclusion chromatography (SEC) profiles showing the complex formation of Binder-82-O32 or the alanine mutant Binder-82-O32-M2 with hIL-4Rα. **(B)** SDS-PAGE analysis of fractions corresponding to the elution peaks in panel **A**. **(C)** SEC-MALS analysis of Binder-82-O32.

A

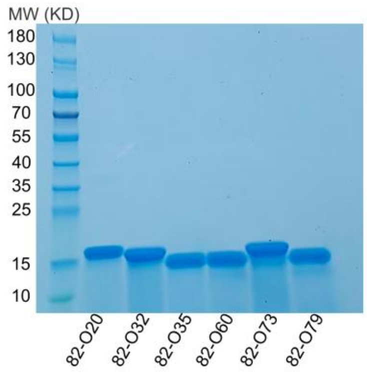

B

| Protein number | Yield (mg/L) |
|---|---|
| 82-O20 | 28.8153 |
| 82-O32 | 60.4685 |
| 82-O35 | 49.6005 |
| 82-O60 | 55.566 |
| 82-O73 | 34.8524 |
| 82-O79 | 45.2767 |

C

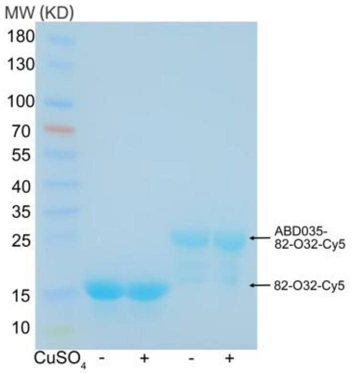

**Supplementary Fig. 5. Yields of hIL-4Rα mini-protein antagonists. (A)** SDS-PAGE analysis of mini-protein binders (5 µg each) following Ni-NTA affinity purification. **(B)** Yield of mini-protein binders from 1 L of expression culture. Protein concentration was determined by measuring absorbance at 280 nm using a multimode microplate reader. **(C)** SDS-PAGE analysis of Binder-82-O32-43C-Cy5 and ABD035-Binder-82-O32-Cy5 under oxidizing (copper sulfate) and non-oxidizing conditions. The gel in **A** was stained with Coomassie Blue; the gel in **C** was left unstained.

**Supplementary Table 1 Data collection and refinement statistics.**

| | Binder82-O32-M2 |
|---|---|
| **Data collection** | |
| Space group | P 1 $2_1$ 1 |
| Cell dimensions | |
| *a*, *b*, *c* (Å) | 48.5, 37.2, 123.1 |
| a, b, g (°) | 90, 95.5, 90 |
| Resolution (Å) | 27.52-3.5 (3.625-3.5) |
| $R_{\text{merge}}$ | 0.175 (0.435) |
| *I* / s*I* | 2.87 (2.65) |
| Completeness (%) | 91.56 (96.11) |
| Redundancy | 2.8 (3.1) |
| **Refinement** | |
| Resolution (Å) | 27.52-3.5 (3.625-3.5) |
| No. reflections | |
| Working set | 5307 (543) |
| Test set | 531 (55) |
| Final $R_{\text{work}}$ | 0.3350 (0.3294) |
| Final $R_{\text{free}}$ | 0.3680 (0.3322) |
| No. atoms | |
| Protein | 938 |
| Ligand/ion | 0 |
| Water | 51 |
| Average B-factors | 53.64 |
| Protein | 54.92 |

| | |
|---|---|
| Water | 30.15 |
| R.m.s. deviations | |
| Bond lengths (Å) | 0.008 |
| Bond angles (°) | 1.12 |
| Ramachandran plot | |
| Favored (%) | 89.92 |
| Additionally (%) | 10.08 |
| Outliers (%) | 0 |
| PDB ID | 24WD |

Crystal structure solved by analyzing data obtained from diffraction of one crystal.

*Values in parentheses are for highest-resolution shell.

**Supplementary Table 2 Amino acid sequence of mini-protein**

| Anti hIL-4Rα mini-protein sequence and response value of BLI. | | |
|---|---|---|
| **Binder No.** | **Sequence** | **BLI-Response** |
| **1** | MgKAAAYQAANAQAGKELKAATAEYTAQVDALNAQGQAALAAGDEAAAQAAADAARKAREALRKTELAIGEKVKAANAAggsHHHHHH | 0.0537 |
| **2** | MgLAEDNRKRGLATIKDAEDQLAKVEARVAEALAAAADDPEKQAVARAYLETAKKEAEKLKAKGKELIAggHHHHHH | 0.0446 |
| **3** | MgEEAKKEAAREVLVSYGKKAKEEIEKAGEELKDKPREEVRKKAKEIWKKYNEKALEEVAKILAggsHHHHHH | 0.0308 |
| **4** | MgAAAAIAAARAKYGAKYLELQKEAGEKIKKAAEEARKKMEEARAEAEAKAAEAADDPEAQRKAWEEAAKKQEEAREATRKLEKENNEKLKALGDELKAEVAAIEggsHHHHHH | 0.0066 |
| **5** | MgEKARELGKKTRQKMLEIESKANKEVGETATSKGLSGEEIAKLGTEALKKNTEEQEKLKEETKKEIERLLAggsHHHHHH | 0.0054 |
| **6** | MgAAQAEVDAWRDAALARVKARMAELLARRQARARELGKRRGASPEEIAAALAEAERWAEESLAAETAAVDKRAEALLAARggsHHHHHH | 0.0677 |
| **7** | METYEKVNAIGKEAKDKADELGEKSRALLEAGSTPEEVLATYQEARDYEKEKHAEATALVEAAKggsHHHHHH | 0.0239 |
| **8** | MKEEALKLLKETEERVEKMEKEFKKLGEEDPRLQPVAKAAEKGAKIAKERIEKAKKAIELDPEAGLEAAKYVKEVVESIEKELKESVKLLggsHHHHHH | 0.0533 |
| **9** | MgDEAEKVIKDAKEAIDAATQKLIDTTKEAAKMGAEDPALAPEARAKYLAARKEYNATYAAASAKAAKAGPEALKAVHDIEKENAKKISAAIEAEVggsHHHHHH | 0.0485 |
| **10** | MgKAEKDIEAIKKGLELQKKKVKATLLARAEAAAAKDPAKGALAYAEAKKRAEETEKFLEEAAEKQMEYLKQAggsHHHHHH | 0.0412 |
| **11** | MgAEEEELAAKEAESAAIRAESQKILDAEKAAAEAIRRASLEGPALLPEAEAALAALKATAAEAAAAHPRNAAQAANAVARAEARLARMRAGLAARAggsHHHHHH | 0.0432 |
| **12** | MTAEDWVTAAKATFDNYVKKAKELAEKAEKGTAEEAARNYERAKRVLESLKTDSDKLIKMAEEASPEAAKEVKKYADEAKKKVEEYVKEAKKAAERKKKEggsHHHHHH | 0.0376 |
| **13** | MgAEAQRLALQQQALAASAEAYKKMKELGEEAA | 0.0847 |

| | | |
|---|---|---|
| | KLRKEALEAYKKDPEEGLKLREKALEKDKEMKKVHAEENKKVAALRAQAAggsHHHHHH | |
| **14** | MgEEAKGAAYLAAQKKAGEEIKKLADETAAKIAAEIAKAKENASDPAAVAAAAEAAKAARDALKKEEKAISDELKAggsHHHHHH | 0.043 |
| **15** | MgAAVAELNAQAAAYQKGLDNTKEIGKAQLAKLKERTAAAIAKAREKYAGLTPEEVQAVIEAAEAEGKEKIAEEQKANDKAISYLQAQRDKYAAAAKEAAggsHHHHHH | 0.0369 |
| **16** | MgDATYKAVRAALKEAKKAAIAGDEENAKKYFQKALDLAAVNPEMQAQTQKRIDAIKKEAAKSRAAKAggsHHHHHH | 0.0782 |
| **17** | MgKIAEVRAAMHKKADAALAAIVAAGNARAAAAPPSQGAAVLAATRAERKAAQAAYSKESLAAIKAAEAggsHHHHHH | 0.0319 |
| **18** | MgGAAVMQAARKAVLDAAKEALLASPEEAPAKIAATAATAKAAAAAAPTPEAGAQLLTVGNAAIEEVKQSQKARLggsHHHHHH | 0.0375 |
| **19** | MgISPEAKAALQKGKEESDAVLQKGVDEANKVAAANPGAEGKEKAKAVRDKARKEAIAIDKEYAAKAQEIEggsHHHHHH | 0.0322 |
| **20** | MgVQAQLQALHDAYTAAANAAVAKANAAAAIAAKSPEIAPLAAAFGKALRKTLAKGSSELPAAQAALDKLKAFAKEAPADVAPYAKAAAAEAEAAIASAQAASAIKAQADALAAAARAggsHHHHHH | 0.0516 |
| **21** | MgEENLRNAQAFLEGAKQTAANGEKAIAKAKALAKERAAKAEAAGDPELAKAILAQGEERAAAIKADEAKGQAYLQQRAQEFLAAggsHHHHHH | 0.0362 |
| **22** | MgNKADEAAANAAAAAAKAKAAEEKAREALAVSPEVARLYADAARAYLKAAEYYRSASLAIKAFPEKADELWATWTAKADAAAAAAEAAEAAGEALLAAggsHHHHHH | 0.109 |
| **23** | MgSLKEKAKELLEEAKKLAEKYAKEAAKENPIVGKAVLTETEDAIKSIEEVPSTLPSAVKNRFDILQKRVAHFPVAQKGLEEIQKLLVEAAKLYQEAggsHHHHHH | 0.0533 |
| **24** | MgAVEKQKEIDAKSKAIEDEIAEKIAAANAEARAIAEAGTPEEAKAAGEAAKALRAELLAERKKKMLELHALAQQAIAEVRAggsHHHHHH | 0.0503 |
| **25** | MEEIEKAREEATKYLTEAKTAKPEEIGKLARKAIKALERAANAAAGLTSDPEVRKQIAEIADKASREIYKLAQKAAASEDPEEAKKYITEALEVAEKALEKLKELggsHHHHHH | 0.0422 |
| **26** | MgEEKAKEQAKLQEAGKEIKEVEEKTYTQAKEEAQKLYDAGAKPEEIIAAIKKAAEETIAAAKKYPILYPS | 0.036 |

| | | |
|---|---|---|
| | YKTHSQKHYERAKERVEKAAggsHHHHHH | |
| **27** | MgKEELLEKALAKKKEYRKKILDAQKEYTKKSAEIAKKYNSDPKLREAKLKEVTDEADKELEKLKEEGNKELEKLLKEAEggsHHHHHH | 0.0318 |
| **28** | MgGEEKLKEVKEKGKEIVEEAKKAKEEVVAAAKAAIALSPEEARKALSEAAKKQYELQKKIAEKLGELNKLAAEAAKESSPELAKKIGEAARKTREETLKELKEMNKAFAEEAKKALAKAREQEAAAAggsHHHHHH | 0.0713 |
| **29** | MgEKIKEEAKKLREEARKKLKALVAEGHKAVAAADKVGPEARLAVGTAYAKKLEEKSKELYEEIEKKVKELKEKggsHHHHHH | 0.0626 |
| **30** | MgDEELKKLAESYEARAKAAKANIEKEAAKLEARTAAAVAKAEAAGGTPAQIEAVRAQGAAKIAENKAFYTKIAEEEAAKAKEYRAKLggsHHHHHH | 0.055 |
| **31** | MgSANEKAAEESKKVFDKKIKEKLAEAEKSEKLAEEHASDPELAAKYRARAKEYRERAEALKKDQEKGVAYLKSRggsHHHHHH | 0.0476 |
| **32** | MgGNEQLAQAALEDGKKAEAKAKELEAKGKVLAKIDPKEGLRALQLGKRYKAEAEFLKKKAEELKggsHHHHHH | 0.0404 |
| **33** | MgAQEKRKEAVKLLKKAVEEASKIWEDAAERIKKEAAELPELTQAYKDAVKKAIEAAKKHAEEAAKAALAGDVEAAAKALRALREEVRAEAKKMLAALAAIAKALPKAAEIAREASAEVEKALDIVHQAAIEAGQLIAAggsHHHHHH | 0.0352 |
| **34** | MgAEAEKLAAKLLEQAKAHNAKAAARREARAIAAGHDPAAARAAAAKHLAEALAIEKKKVDAYLEQLKAAAggsHHHHHH | 0.0457 |
| **35** | MgEESILEKIAKRATPEQLLATLENARKALEEAKKLAKTTPEKAKEAGKLAGRAAATATEAAALAGIKGALTPEEVLKINKEAVEALLEVYKENPELVKEDVLRALRRSAKIMEEAAKVDPKNKEAFEKAAKLFEEAIKEVggsHHHHHH | 0.0425 |
| **36** | MKELAKKKLEQAKKLLEKMKAAAEKAKALLEAIRANPDAAPELRKAAGKAAREFMKYLLATAGAASAALRYLGEPAEVSDKVVADARKAVDAVYETTNAALADPAAAPAAIKTIEDALKKAEEAVKYAEKLLggsHHHHHH | 0.3818 |
| **37** | MgAAAAKAQAKKELAKGEELLKKAEELLKEAERALKEDPEKALKYAAEAGEAVREAQKHYEKAAQYYALAGDYATAKAVLEEGKEKAEKIRERILKVGKEAEERLAAggsHHHHHH | 0.0586 |
| **38** | MgEEEARARAERDIAAVEAAREAAKKRIAARAEERIKRAREKAAALGDPELAEQAEAEAKEKEEESKAAE | 0.0631 |

| | | |
|---|---|---|
| | DERAEKQIAYLRQHggsHHHHHH | |
| **39** | MgDAAAQMAAVRKEAGEEVKKLGEETAKKIEELIKKAKASTPEEAAKYAAEAKEIRAKAKEEEKRIDEKSAAAIRAIEAAAKggsHHHHHH | 0.0665 |
| **40** | MgSAVLKAIEEATKKAIEAAHAFKARFEELAKEGRAAEALAEVKAQAEAEAAKATPEARAGIMAKIAAGDAAANPSQAMRYYIKALEGAISSRAGAEARAggsHHHHHH | 0.0545 |
| **41** | MgALEKAKEVAKKAREEMLEKVKALYDEAAKAIKAADKEGPEAYKKTSIEETEKIKKAAEELKKKAEEIAEKALKKAEEELKggsHHHHHH | 0.0656 |
| **42** | MgKEELEEAKKITEKYEKSAKEAEEKLKKKMEEYKKKAEEAEKLDPALAKQYKAKADSLQKELEAVKKRRKENVELLKKLLEEggsHHHHHH | 0.0599 |
| **43** | MgSAAAEVAALQAAIKQTNAIYKTRADEALKLAKAAKDPKAKADYEARAKALLDDAKKGEAYLTARAAAALAAggsHHHHHH | 0.0896 |
| **44** | MgSPEEIIKNLKKKREEYLAKAKEYKAKAKADPEKAAEYEAKAKLYEEAAKNTEFHIEIEEAALLAKKAEEEGDAEKAEEAAKKLEELAKKASPKLGKALKERAKRARAFAELLKKKggsHHHHHH | 0.0059 |
| **45** | MgSAALEEKKKIYQELAEKSYQKEKEAAEKIRAAVEEARKKAAEDPSKAAEYGEAARKEREKLKAEVVKFQQELLKEYEAKTAKLDAEggsHHHHHH | 0.0834 |
| **46** | MgEELKKAAEAAKAAAAAAHATALALHKKSGEARSAAAKETDPAKAAALLAEADAYFAAAKAAEDAAGAARHAARALQAQVEAggsHHHHHH | 0.39177 |
| **47** | MgAAEAAVEAAIAAAKDWGKKAVEYAAKVDPALAEKIKAETAALIAAAKAQPDLAARQAAARAATKEMEKKFNEAYGLYQLKKQKELEALAggsHHHHHH | 0.0818 |
| **48** | MgAAAAAAAAAARARIRAAQKAGAAALNKTAEASKSLSPEEQQKAIAEEAKKHDAEIEALYAQAQAEIQAVLEAggsHHHHHH | 0.0794 |
| **49** | MMKEGIEKRFEKLLELAKSEEAKEIAEKGKKEALELYEKHQELGKKAQEYSADPSTAAEAEKARKEAAETEKKLGETIKKTGEEAGKIILEEKAKggsHHHHHH | 0.0186 |
| **50** | MgEEAVEQARALAEQAVASAEAAIAKLTKLRDEYAAKAKEAAKISPELAKQYQAKADSLQKELDAVKARKEENLAKLEEIEAAggsHHHHHH | 0.0711 |
| **51** | MgDEAARYLASAEAVKAGQEKVLAKLKAKGEAVAKEKAADPAAAAQAKAHYEKALKEEKAKYDKRIAELEALAAEARAAAggsHHHHHH | 0.0819 |
| **52** | MgSLAEQQKIYEQSQALGKEARALEDEAGALLAAALTSTPAETLALAAAAAAKLVAAGELHAQSHRLAG | 0.1076 |

| | | |
|---|---|---|
| | DEAAAAATLARALARAARLVAAALAAVAggsHHHHHH | |
| **53** | MgLSAAEREARIAADFAAIEEGIKRATARLEEIVEKRKAKAEKLAAELAKEITPEEAQALLEAAKAQAEEDREINLKFIKEAAEKRKAYVKAAggsHHHHHH | 0.536 |
| **54** | MgAEEIAKQFFAGAENTRKIAEKRREELKALAEELKDNPEAAADALKRAEEDYAFHEEQAKYLEDKGKEALAEggsHHHHHH | 0.0988 |
| **55** | MgAAAAAEAAAAAAQYKAAAAQAKANRAKKAAAYDAAAAAAAAVSPAAAAAVNARKAGADADYDKGIAYLTAQAEKFAAAggsHHHHHH | -0.0398 |
| **56** | MgKKLAEEALKEAEEKKKELEEDIKKLKASTEALAAKTIAEYPETGPKIAAEAKANTEKQVAKLKERAKEIVEKAKEFAEggsHHHHHH | 0.0065 |
| **57** | MgGAAAYEALQAQARAKIAEIAKKTDARIQAIKEQGEAEAQALLAAGDLEAAKAVAAAARAAMRAAAKDQAAESQKVADTLAAEARAAEAAAAggsHHHHHH | 0.0101 |
| **58** | MgAEWFAEAQAFLAAAEASKAKYDKAIKDRLARAKELAAKAPDPADAAAVLADAEEAGKVIKKDQEKSVAYLKAKAAEFAAKGLAAggsHHHHHH | 0.022 |
| **59** | MgSEALKVQKEIKEKRDKMRAKFKELLKTVNEAALAADRRDPAAAKKLQALSQELIEAEKEASNKFKKAGELAKEDPEKGKELADEAEKEAEKKLEEISKAAAAVAAggsHHHHHH | 0.0047 |
| **60** | MgGVEKVKEVAKEARAKAEEYRAKAEKTTPGEAQAWNRGAVEAEDIARGADYVISRAPLSSDETVAQAAENLRKALEKAEERAEARAARVRAAEEAAggsHHHHHH | 0.1164 |
| **61** | MgSAEEIVKQAIEAAKKWAKKAAEYAAKVDPALAKKIEEETKEKLKKAEEEKDPAEAQRIAREAAKEAEKKLNEAYGALRRKEEAARKAAEggsHHHHHH | 0.2277 |
| **62** | MgSVYEEAEKAYEEALEFNKKVAAAAQERAKKNPEDGQKMLDQAEKERKDIEKRRKENEERLELLKAAggsHHHHHH | -0.0444 |
| **63** | MgSEELAQQAQKLREEYRKKMKEAQKLLAAAVEATRTKSPAEGKKYLAEAEKVFKEALEIMKEAAKVAAKVDPEVAQLYLEGLETDTKAFEYSKALINAEIAREEARRggsHHHHHH | -0.0433 |
| **64** | MgAAAQLEKDGKAAIENLKANTEKKVAAAKEKAAKSTPEEAAKELARAERYKESYEKDTEKAKKAWQARIDAAKAAEAAAggsHHHHHH | -0.0275 |
| **65** | MgAEAAKKIKELAERTVKEIREAAAKGLAAGKPELAVEESRKIAVEGHNEIGRIATEAIAKGTPEEAKKVIEAAREGLKAIGEAVKEAREAAARLAAALAggsHHHH | 0.0146 |

| | | |
|---|---|---|
| | HH | |
| **66** | MgAAELEEARREAAEEQRALREEVREKEKAAQEQYGALRKAGQADPSKAAEYNAKATAIVKEALEFGKETGEKIRAIIEEVEALAggsHHHHHH | 0.0159 |
| **67** | MgDKEQLVEEAEKQGKEDLAIEKADVEKGIAYVQAIRDAHAGDPEIAAYAAKGLEQAKKNGEKRIAKVEKRHKEAIEKLKELggsHHHHHH | -0.0188 |
| **68** | MgSEEELRARSEALAKLSQAVEDANGAAVQAAQDTAKDPAAAPAAKAAIAKARALGAELPEMKSATDKTIARVEKGLAAALAggsHHHHHH | 0.0079 |
| **69** | MgSEAVEQALANAKGYLEGKKRHEKVMAEKAAAATSPERKKAYAESLKTETEIDDKAAADALAKAKALLAAAggsHHHHHH | -0.0389 |
| **70** | MgAGVEKAKERKKERLKRIEEGEKLVEESSPEKKLEIAELARQIVENNEAGAKLTTAAREANLKRLTADEEAYLAEAAASTDPAVKAELKAKAAAVAKAREDSDALLTAEIDGQKEVAEKAKEVAKKAEELAKAggsHHHHHH | -0.0194 |
| **71** | MgAAAAAAEAARRAAGGAEVQKLLAEMGKLLEKTTKKMQEVRDLALKAGEASEENVEEGEKLAEKAFKKAEEVVKDTKEEAKKLADKAKASTALTPEEKEALVKGAELLVKEVEKYAERLKEAVSKKIELTKAggsHHHHHH | -0.0033 |
| **72** | MgSLAEEEAILAEAQKKFNEAAKEFNKKFKEAGEKAKAALEKAHAAAEAGDKKAMDAALAEAKKALEEMRKAAKEAAEKAKEIGKLSPKAAEYAKNFAKQVKNQVDKFGKRVEAAMKAIRAILAAKggsHHHHHH | 0.0022 |
| **73** | MELKERTVKGTKLAAEKTVKRKERLDAAAAKNPGEAGALLKKAAEAETLADAYEIAAGVLNAAASAQALDGSGPEITKAGAEVLAKTIPEAAPIAEAVKKAGAGTAEELKAAAQEFMAKAKELAAQATAYRSAAVATAAAggsHHHHHH | 0.0674 |
| **74** | MgAERRERLLKEAEELAKKATEERTALVKEINANLTAAYAAGNAGLEALLAAGAPREEAARLARRVREALRLVQSTPEKAKEGVELAREVAREAAALGYPEAAALFEAAAEALAKAAEARAKSEALAKEELERTRRLLEEAEggsHHHHHH | 0.1132 |
| **75** | MgEIEKLLAEAEELVKKALKIVVDAGKKAGVEELLKEAAKKAEELFKEALKLYEEGAKSESPEEARKLRKEGNKKFRKADKEIGDAVREAQKKVTPEQAKILTDALNKASKLFIEAEKLANEAVKLAAKEKYEKLKggsHHHHHH | -0.0244 |
| **76** | MgAQAIAEEEAARQAHGAQVQALGKAQWAEVQKAAAQARAVTAEAAALAAADPSVAPLAAELGRAMR | 0.0305 |

| | | |
|---|---|---|
| | AAVAALEKSAGYRTAAANAPPAEAAAYNAKAAAYKAAADAAWAKAQALLAQVRAggsHHHHHH | |
| **77** | MLEENLRILRELREVAEEVAKATAADPAAAAAFQRALDLIDEGERRAEEALEEGDEEKATLIAEEAARRAASAVLEAMASAVAYYASKVLGFAVGQKLAAEIIKKGEEAVKNLELALKYAREARELVVELAKKAggsHHHHHH | 0.0112 |
| **78** | MgSLLEELKKQAEIEKARQRAKRYYESADEIGERSDKLAEAALKAGDAVGAMIALAGAAHAYASAAVKLAELAKKAKELGDEETAKEAIELAKELLEKAKEKVKEAEKLGKEHPETAADAKIAKESFEELAKKAEELIKKAKggsHHHHHH | -0.0134 |
| **79** | MgAAAAAEALAEQAGKDEIAAYAKAQKKLDEATTAAKELLQALAEKALADPRYAAAAKAAADAYKAVNAGFKKLLTDPDAGVKAVAAAMDTVVAAAAGELAAEAKAVADEVTAKAKEAAEAIKQAKAEAAAARAAVEAARAggsHHHHHH | -0.0061 |
| **80** | MgAAAAAALERAALAKAAAEQRAAAYQAEIDRHTAAYEEGKKLAEALRKADPSLTPEEIAAKLIEAAKKSTDPAKRLAYSEGAVYAENPEKAKEGLARHVARHQARIAEARAAAAAVAAEAAAAIAAAggsHHHHHH | -0.0189 |
| **81** | MgSAAAEQAAKIAADADTATTNLNAQAKKAEAAGQAYAAKGEAATPEEAAKYYAKAARAYRKAADYYNSASLFANQKSKQLAALGDAATAAALAAKAAEYNANAEKAKAKAEEAEAKAKELKAggsHHHHHH | 0.0943 |
| **82** | MgVEAWREEARRLIREGRRELLAVFREQAKLITEAAKKAEKSTAEEAKKILEEKKKEVTERITAVTARIRERLEAAAERAPPEVAAVLRAGAAQLAEIGDTELAALLKGIDDIEERVVAHLKggsHHHHHH | 0.2948 |
| **83** | MgLTEKAAEALQKARDATKEAELEVNKAYGIISTAAQEAKELNPEAAKELQKLKDELIKEYQEAKKKIQEEYAKAAAAAQAGSTLEEVQAWADKAIDEAKKLKEKAKKLEEEAKKILEKTKKEAggsHHHHHH | 0.0085 |
| **84** | MgALEAALKLMEKALKIAKEAAEIVSFDPALKALMERAIKIIEDAMAKAKEYKAAGDLEAAKKAAEAGLKRAVASTLEALSGAVRALVSRTLGPEAGVAAAKEIYALAKEAVANPEKAAELLEEAYRRVAEAAKLAggsHHHHHH | 0.0248 |
| **85** | MgAAAAAAAAAEAAKAAAKAKSDAALAALTAATAAGTATVVGLATEGIAELARAAAEGPEALREAAARVAAAIRAAIAAHRATANALSTEMMTAASKAGPDQYAAANAKVGELEKTYAATASAAIQALVAAggsHHHHHH | 0.0079 |
| **86** | MgEAAAEEQAAKEQQVINDAGKKAADAIEKIGEA | 0.0023 |

| | | |
|---|---|---|
| | AQRADADPSLAPKEAEVAKKEADAIVQAAKDAAAALTNKEWAKNFALSLATALEKLSKRVARSPYNQAIADTLKAASKELKALAKALggsHHHHHH | |
| **87** | MgDRVERVKEAVLEKAEEIAKKDPKEGRAMRPALVRVAYHSTLALLAAERAGETERAEALAERALAAETREEAIAAAAELGKLAHELLIKVGEEAPEILKEAFERAKEAEKEITEKIEKIAKEKggsHHHHHH | 0.1198 |
| **88** | MgSAAAALAAEQADRAAKAAAAAAANKAFVDGEKKVADAMNKLVAAAKEAAAKSPQAEAVGKLARDTYDVYKTKGKEAAAALAAARTAGLAGDAATAAATYAAAVATIDATYDDVLKAIDAALAQVKAIAAAggsHHHHHH | 0.0057 |
| **89** | MgVEAQAKTVLEQMKKAVELVKEVAKKAPAFAPTAKLVEAALKFHQAQIELDPEKGAKAAGQALAVMAVEAAKSAMRALVQWAVETGEYDEAVEILTKMSEELTELVKKADPSLADKTAAATKAAVDEAKELAAKAKAAggsHHHHHH | 0.015 |
| **90** | MgKEAVEKAKKELLEALKKLVETLKKATELLAKHSTPSQAKAAQYDLDKAKEAAEKAKEAAKKSPVEAAKEARKAATYADGAGGLALKVPETAELGKELKEEAEKIGKLAQELVKAAggsHHHHHH | 0.0062 |
| **91** | MgELAKVAKEAAKEFEKIAKKAKELAKKLEELAKKTSPEAAQHLLRAASLARELADIAKRAAENLRAAAAGSPSDAPVAAAYMTSAAEAFKKLAKKLEEELKKFKATGEDNPEAEEAIKEAIKLAKKAAEIAKKVAEALKAggsHHHHHH | 0.028 |
| **92** | MgSEFEELQKKREELGKVVAEANKEAGELSKEMAEKFAKINAELTALAKAGTPEEAKIIGEGERHLRLALLYAENSNTPENVEKAKKHIEEAKKYAEASKSPEAKKTIEKGVEAAKEYLEKMKEINDKLNEARKKSEELggsHHHHHH | -0.04 |
| **93** | MgGKEVLKKALEEAKKLAEKVKELAKKVEEYGMKGIGTELKKLAKRLVETVEGAVKNPEGIPLAIENLKRIEEAAKEIEKKAQYVVELAKKDSPEKAEKAKKALEEAKKAAEILKKAAKEAIKAFKKVLKELggsHHHHHH | 0.0351 |
| **94** | MgAAAAEAAALAAALKQAQLDDKTTRDALEQAVKDAAEELAKIMKEAIEKEGLPEEAREAAKKAVKLIRKAAATPSKADEYLKEAEKEVKKLAELASEEYAKKIEEAFKKVKDAIDALKAATAAGTAKITALSAQIVAALEAggsHHHHHH | 0.4043 |
| **95** | MgAKQAEIEAAGKELKAARDAARKELKAINKKIAEASKELARNGNVEMQEEAQKIINEYGKKIKEAEEKAGEYIEKAVKDPSNIKEYLAKAKEEFDKAVEIAKKLL | -0.0026 |

| | | |
|---|---|---|
| | EEVKKMAEEAKKKggsHHHHHH | |
| **96** | MEEAINGAKESAKKHIEKVKEAAKKAIENAKKAKNVEEVKAVAEELRKAFRDIQAEHNKDDQKIGAKIRTLSEEDAAKAQAVLNEESGKFIDTMREEGGKAIDTLLEMAAKLEAEEggsHHHHHH | 0.016 |
| **97** | MgVTREEVEALAAEAKKKIKEAVDKERDAGLAHVAKIKAGATPEEIGAAYAAHQKAGAESEKTIGDALQALRDTALEAYRQALADPRYAPVAELARQEYLDARQTYKDTRSKANAEIQAALAAggsHHHHHH | 0.0158 |
| **98** | MgAAAELLAQAKEKAAKGEEAAKKAIDLAKAAAAALTAAAPQPPEVIQALYIAESKKVLDAAGKAMALMREAILLAKAAGPAAAAEMQAKYDAVKKKLADARNALAATYQAAYAAAQAAggsHHHHHH | 0.0756 |
| **99** | MEAIKATEKELAALAAAAEALAATLKAAGVPAKYSDAFLAAGQKAVANLQKRAKKAVEGALALPDPEEAKKLLAELAKQLKEATATLIADMTAGAAYIAAKYAAGKAAAAEAAALAggsHHHHHH | 0.2875 |
| **100** | MgVEEALKAAGEAARARYRAENAKAYAEALAAAAKAMAAADPALTGAAATAAADYAKQAIANAADPEKQKKYMAEANTAYATALAALSKADPAKAAAILKAEADAYKAASEKVYKEVNAAAEKELAAIKAggsHHHHHH | unexpressed |

| **Partial diffusion optimized sequences for Binder-36 and Binder-82** | | |
|---|---|---|
| **Binder No.** | **Sequence** | **Relative luciferasence activity (%)** |
| **36-O1** | MEALKKELVEKAEEALAKAEALAKELLALLETILAAGELTPEQAARLRTTGRALLKAAETAVSYIVALYRELGEPQEKQDEVFNTSLTLIQAVYDALNAALADHRQGPALIAAVKALLDYLTKELEKAKELAggsHHHHHH | 39.75114957 |
| **36-O2** | MgREELIKALLEAAKAAVAKLKATQAKVTALANKIAAAATVTDEEKKALGDATRAFLKDAETAMSYILAAYRELGTPQEEIDARFKEHNALIQAVYDAANAALADKSKLPAFIAEVNALLAHVEAQLAEVEKLAggsHHHHHH | 66.06437653 |
| **36-O3** | MgTQARLDELIAKAQAALAAAQATAAEVRALVAELLAAPTATDAQLAALTRATRALIKAFETSMSYILAGFRLLGAPEAEVQALFDQGKALADAVAAALAAFLADRAHAPAFLAAVDALLQFAADQLEKVTAAAggsHHHHHH | 46.83797673 |
| **36-O4** | MgSEATLKAVLAQAEAALARAEALAARTRALAARVAAAPEATDEQLRELGRQVRAFLRAAE | 58.83148497 |

| | | |
|---|---|---|
| | TLLSALVAGYRLLGKSEEEAMALFEKANALINAVWEAAQRALADRAHIPEFVAAVDALLAYLRAALEEVRALAggsHHHHHH | |
| **36-O5** | MgSAERLEALLAAAKAALEELRAAADEVRALVARLLAAPEPTPEQLLALGRALRRFLRLAENLLGLIVGVLRELGVPEEEYLAIAKELLALITATGEAGQAFLRDRSAAPAFLKAVDDTLAALEAALERAEALAggsHHHHHH | 32.25244547 |
| **36-O6** | MgKEALLREQLERARALLARARAAADRLLALVDRVEASGAPSDAELREVGRTARAMLRAAEAAASAIAAVFRLLGKPQSEINAIVANTNALVKAVADLLNKFLADPSTAPEFRAAVLALLDYLEGLLEEAERLAggsHHHHHH | 50.9277793 |
| **36-O7** | MgREALLKEVLKKAEEALEKLKALAEEVKALVERIRAAGTPTLEQLRALREAVRKFIKAAETALSYIVAAYRLLGLPEEEVMAVHNEGLKLINEVYEAAQRALADPAHAPEFLAKLDELIEYIEKKLKEAKKLAggsHHHHHH | 92.85366513 |
| **36-O8** | MgKKEKLKEYIEAAKAALEKAEAAGKEILALVDRILAAGAPSDELLKALGDATRKLLKYAETATSYILAALRLLGVPEEEINKVFNEMNALIQRVYDAANAFLEDPAHAPELRAAVEETTKFLKEGLEKVEKLAggsHHHHHH | 32.9402218 |
| **36-O9** | MgKEALIKELVANAKAQLAKAKALAAELEALAAKVLAAGEVTPELGAAFRAKARAFLKAFEAAMSYILAAYRLLGKPQEEIDALWKKGQELIQKVYDAANAALADPSKIPEFLKAIQALIDYAEAELKKVEKLVggsHHHHHH | 66.42683253 |
| **36-O10** | MgRQALLDEVLAQAKELLAKAKEVAAEIKKLATQLIKAPEVTDELKKELGKKVRELLKYAEAAMSHIIAALRLLGAPQEVLDSRFNKALTLIQAVYDAAQKALADRSHVPDLIAAVDALLDYLEAELAEAEKLAggsHHHHHH | 100 |
| **36-O11** | MgREAAIKELLKQAKEFIKKLEEKAEEVERIAKEIAASPEVDPEKAKELGRKTREVLKLAEAAMSALVAAFRLLGKPEEEIQKYFNEHLKLLQEVYDAAQAAIADRSKVPEFLAKVKELIKYLKELLEEVEKLLggsHHHHHH | 74.3738166 |
| **36-O12** | MgKQKQIEELLKQAKKKVAEMKAHRDRALALVDVLVADPNAPEAVKLELRRTVRAMLKAFESAVSYLIAIFKLLGKSDEYLQAKWNEAKALVQAVYDAANATLADPSAAPALRAALDAAEAYLESLLKEAEALAggsHHHHHH | 7.286989433 |
| **36-O13** | MgTQAELDKLLAAAEEKLKAAEAKAKEALA | 29.19097533 |

| | | |
|---|---|---|
| | LMDEIAADPASPDEVRRELGRKVREFLKEAESLLSYLVAAWRLLGKPQEEVDSLFNTGLQLIKDVYDATNAALADPAKLPDARAAVQKLLDFLKEQLKEVKAAAggsHHHHHH | |
| **36-O14** | MEALLQEVLAQAEELLKKAEEELKKTIELATKLLDDPNAPDEVKKELGKTTRKYLKYAEALASAVAAGLRLLGEPEEEVMAVIKTTKALIDEVYAATNAALADRSKIPAFLAALEKLKAYLEKQLEKVKAAAggsHHHHHH | 28.4697026 |
| **36-O15** | MgKQAAIDELLAQAKAFLAKLKAKAAEVLALIDKIAADPNATEEELKALAKAVRAFLKALESLQGQLVALMRLLGWPEEKIQQVFDDFNALITAVYNAANEAVADPAHLPALAAAIKAALTKAETALAEAEKAAggsHHHHHH | 29.00496747 |
| **36-O16** | MEATLERVLKQARELKKKLEALKEKVVGLAKALAANPNPTDEQLRELGKLARAYRKAFESLTSALAAARGLLGAPPEVQQALVDKAKEMSAAIYEATQAFIADRSNAPVLIKTIEEATKYILGQLEEVEKAAggsHHHHHH | 16.2076285 |
| **36-O17** | MgREAAIRAALAAAKAALAAAAAARARLRALLAELAANPDASPEFRKELGATYRAWLKAAETLNGLIAGARRLLGAPLAEVVARSQAILALITAAGEAFRAAVEDPAAIPEFLALADQVAAALAAALAEVAAAAggsHHHHHH | 24.18627043 |
| **36-O18** | MgRQAQIDALVAQARAFLARAEALRDELLALVDKIAADPAAPDALKRELGRLARELIKAAEALTSAITAALRELGAPQEEVDAVFNTGLELINALYAALQKLLADPSHAPDFRAAVDALMAYLRSQLERVEALVggsHHHHHH | 38.2003195 |
| **36-O19** | MgSEALAKELLAQAKELLAALKAARAEVLALAARIAASPNPSDEDKKALIAATRKVLKLAETLVSKAVAALRELGASEEEYMGLWKEMQVLIQAVADATQAAVADRSKVPELVAAVDALLAHAEAVVAKVEKALggsHHHHHH | 30.89699543 |
| **36-O20** | MgTREQIEALVAQAKAFLAKAKAAAAAIKALVDKLVAAGTPTEEQLKELQTLTRALLKAFEASMSYILRAFELLGAPAARVQALFDAHLALVDAVYNTANAFLADLSNAPAFLAAVDALLAHAEAQLAELEKALggsHHHHHH | 14.28014967 |
| **36-O21** | MQKELDKVLKQAKKLLEKAKKLAKEVLAKIDRIEASGAPSEAELRELGELTRKFLKALESLASAIAAGYLLLGKSQEEANAIITTTNAAVDRIYALANAAIADPAHLPAFRAAIEEALKTAEEQLKKVEEAAggsHHHHHH | 21.97824803 |

| | | |
|---|---|---|
| **36-O22** | MgRQAEIDRQVALAKELLAKFLAARDATLALIDQIAADPNPSDALLKALGTETRKMLKYAEAATSAITAALRLLGAPQETIDAVFNTTLALVQAVYDAAQAAIADPSQLPALRAAIDALTTHLTAQLATLEALAggsHHHHHH | 25.46119 |
| **36-O23** | MgRRETLESLLAQARTKLAEAEELAKKAKALLEAIKKEGAENVALKKEFGKAARAFLKAAESSMSYILAAMRLIGLPDEEIMARFKEFNKLVDAVYNAANAALEDLSKAPEFIKAADALLAFIKAELERVEALAggsHHHHHH | 19.21351513 |
| **36-O24** | MgRKELLEKLVAQAKKALAKAEALRAELDALVAQIRADPSAPDEVRRALGETTRAFLKALETATSYVVAAFRELGAPQEEVDAVWNTTKEQVDRIYGLANAAIADPAHLPAFAAATADAIDYITAQLAKVEKLKggsHHHHHH | 37.9744841 |
| **36-O25** | MgRQALIEQLLAQAKALLAKAKAARDKALALMDEIAANPNAPDAKKKELGKAVRAFLKAAEALASAIAGAWRELGASDAEVQGIVDTTRAQIKAVYDAAQKALADPSAIPEFKAKLQTLLDYLAKQLEKVSKAAggsHHHHHH | 27.22454427 |
| **36-O26** | MgRQARLAALLAAAEAALARMEAERDAALALVDRLLAAPEPDPELLRALGRETRAMLKAAETATSYIAGVLREAGAPQSEVTGVVDATKAAIDKVYAAAQALLADRSNAPAFRAAVTDLLARVRAEHARAAALAggsHHHHHH | 33.3286649 |
| **36-O27** | MgKKEQIQKQVDALKEKVKKLEALAKEALALLDKLIAAGKPTPEEKKALGDLVRAALKEYETAVSYYIAALRIAGAPQSEQDSKWNEGLALIDAVYQAAQAAIADPSKAPAFVAALDKLLNKAKEYLKEAEELLggsHHHHHH | 26.84902727 |
| **36-O28** | MgSQAQIDALLAAARAALATAKAHADALRAVVAQMLALGRVDPALAAQARAEVRALLKAAETATGYITAILRLLGAPQSEIDEVFNTTLKHIQAVYDAANAALADPAAAPALLKAVDDLLKYLEAQLEKAEKLAggsHHHHHH | 86.73654727 |
| **36-O29** | MgREALLKELIAQARKALDKMKELKAKILAIVDRAVERGELTPEEGRELGKLVRSYLKAAETATSYIVAAARLLGWPEEELQQIFDTTQTLIQKVYEEANAFIADLSKAPELKAAVTELTDYLAAQLEKVEAAAggsHHHHHH | 92.2117426 |
| **36-O30** | MgSQALLEALLAAARAALATATAAAATALAALARIAAAPAPDPALLRELAAAVRTLLKAAETATSYIVAALELLGAPEEERQRIFTTTQTLIQAVHDAALAIATDRAAAATLAAAVTALTTTLAAQL | 100 |

| | | |
|---|---|---|
| | DRVEALAggsHHHHHH | |
| **36-O31** | MgTEELIKELLERGKELLKKAEELGKEVIELA<br>ERIRASGNPTDEELRELGRLTREFLKAVESLA<br>GVLVAGYKALGKPQEYQDKLFNETLKKIEEV<br>YELAQEAIRDLSKIPEFIKAVKETVEWMKEKL<br>KEVEKLAggsHHHHHH | 50.9343 |
| **36-O32** | MgSEALLRELIERARRALEEAERLAKEIKAKV<br>AELEAAEEVTEAQKRELGELVRRYLRTAESL<br>AGHILGAFRQLGASEEEILSLHKEFTELIEKV<br>YNAANAFLADRSKAPELLAAVEELLKLLRAK<br>LEEVERLAggsHHHHHH | 109.6155109 |
| **36-O33** | MgRQELIKKLLEQAKEALKKLEELAKELLAII<br>DKLIASPTVTEEELKALGDTTRKALKAFETL<br>NSYIVAALRELGAPQEEVDAVFNKALELIKK<br>VYDATNKALADRSKLPELKKAVEELLKFAKE<br>QLEKVEKLAggsHHHHHH | 47.02666667 |
| **36-O34** | MgTQARLDELVAQAKAQLAKAEAAAKELLA<br>LARALLESPEVDEALLAAMRKTLRAFLKAAE<br>AANSYLVAVLRLLGAPEAEQQGVFNTILALIK<br>EVYDAAQRGFADRSHLPELIKAVEKLTTTLK<br>TTLAKLTAAAggsHHHHHH | 76.6269 |
| **36-O35** | MgSKEKAKELLKKAKEALATAKALQAAAIA<br>LLDQIAADPAAPASVLAALGAKLRAFLKAAE<br>TAIGYISGAKRYLGYPLEEVVAKSRELLALLT<br>ATGEAGKAALADPAKRPEFVAKSAELLAALE<br>KELKEIEKELggsHHHHHH | 47.91763333 |
| **36-O36** | MgARALAEELAAQAKALLAKANDLKAKVT<br>ALVDALIANPNQSDEFKKEIGDTTRAYLKTV<br>EATTGAIVAALRELGEPAAKADAVFNTTKALI<br>DAVYAAAQAAIADPSKAPAFKAAIDTLDAYV<br>TAQLAAVEAALggsHHHHHH | 29.19097533 |
| **36-O37** | MgSAALLERLLADARAALAAAEAEARRLLA<br>TLDRLAASPNPTEEELKELGRLFRAFARHAET<br>AMSYITAVFRLLGVPEAEIQARVDELRALLDA<br>GAAAVNRAAEDRAALDAARAQVEAALAHL<br>RAQLAEVEALVggsHHHHHH | 28.4697026 |
| **36-O38** | MgRQALIDTLLANAKAALAKAKALKTTLLA<br>QVAAIAADPAAPEAALQALATTARAFLKASE<br>TLMGYVLAILRELGVPEEEILALFNEHNALIN<br>ATYQALNDFLADRSAAPAFTANVDALLAYLE<br>EKYAEAVALAggsHHHHHH | 29.00496747 |
| **36-O39** | MgTQALLDQQLANAAALQTKMDAAQARVL<br>ALLDEIEALGAPSDALLRELGREVRTFLKYAE<br>AYNGAIIAALRYLGAPQAELDALHQTTLAMI | 16.2076285 |

| | | |
|---|---|---|
| | QAIYEATQRALADPAHLPDLRAAIDALKAYL TAKYAEVAALAggsHHHHHH | |
| **36-O40** | MgTQALIDELLKQAKAALAAATAKRDQILAL LDQIEADPASPQSVLQALATTTRAFLKAAETL TSYIVAIYRLLGAPQEKLDEIYQTNLTLVQEV YEAAQLALADRAKAPLLREKVTALVDTNAA WLEEAEKLAggsHHHHHH | 24.18627043 |
| **36-O41** | MgSAALAAALLAQAEALLAAAQATYERLRA LAARIAAAPEVTDEQLRELGRLTRALLRHAE ALTSALVAAARELGFPQSVLDDIETTTKKLIE EVYAATQRALADPAHLPELLAEVDKLLAYLQ GQLDKVRAAAggsHHHHHH | 38.2003195 |
| **36-O42** | MgSSALLTRLLARAEALLAEARALAAEARRL VAALLADPAPTDEQRRALGRVVRALLRTAET LVSNLVAAYRELGKPQEEQQALLDTARALID DVWSAANAFLADLAAAPAFLAAVDDALAFL EAQVAKVRALVggsHHHHHH | 30.89699543 |
| **36-O43** | MgSEELLKELLKQYIKAEQKAIDTAKEAIAKV EALLKSGNPTDEEKRELGKLTRELLKAIETAT SYRVAVLRLLGKPQEEIDKVFKESQEKIEKLY QATQDFLADPSHAPELLALLKEIIKEAEKWK KEAEKLLggsHHHHHH | 14.28014967 |
| **36-O44** | MgSEALLAQLLAQAEALKKKADALRDEVRA AVAALVAAGAPSDEQKRALGKAVRAFLKAA EALASAQTAALRLLGASEEEVQGRYKEALAL IQAVWDAAQAFLADLSQAPAFLAALDALAA YLDSLLEVVKARAggsHHHHHH | 21.97824803 |
| **36-O45** | MgTQAQLDTLLAQAQAALDKAKAALAKLTA TVQQYIASPEPDPALLKAIGEETRAFLKAAET ATSYIVAAFRLLGTPQSELNEIWNTTKTLIDAT YQAVLAIPEDRAAAQTLIAAAQALLDYQQEK LAEAKAKAggsHHHHHH | 25.46119 |
| **36-O46** | MEELIKEVLENAKELLKKAEELKKEVLALVD KILAAPTPTEEDLRELGKKVREFLKYAESATS SIVAGYKLLGVPQEELEKIFNKTLELINKVYQ AANDFLADRSAAPKLKSNVEKLLKYLKEELE KVEKLVggsHHHHHH | 19.21351513 |
| **36-O47** | MgTQEKLKELIKKAKELLDKAKALADEAIAL AQQLLDDPNAPDSVKKALRKTVRAFLKAAE AAMGYILAGFRLLGKPEEELQKIHNEHLALI DAVWNAANAALADRAHIPDFIAAVKDLLDF LEKQLKEVEKLAggsHHHHHH | 37.9744841 |
| **36-O48** | MgRQEAIQAALAKAEAALEALTAAADALKA LVAKYVASPEKTDEMVQAIMTQTRALLKAA | 27.22454427 |

| | | |
|---|---|---|
| | EALAGWLSRVFRLLGAPEERIQQIVADTDALI DAVYNAANEAGKDRSKAPALVAAVDALVDY LAALLEEARALAggsHHHHHH | |
| **36-O49** | MEKLAKELLKQAEKALAAMKAAIDAALKLI DQLIADPNPSDELLKALGDAVRAAMKAFETF TSYIVAIFRLKGAPQEKLNEIFNSNKELIDKLY KAANAALADRSKLPELAKEVKNLQKYADEQ LKEAKKALggsHHHHHH | 33.3286649 |
| **36-O50** | MEKLLKELVEKAKEFLEKAKEYAKKAKDLL EKRIADPANAEKYDKEIKEAVRKMLKYSEAA LSYIIAAKRLLGKPEEEVMALFKEHLKLVND VYETAQAAIADPSKAPEFNAAVDKLLNFLKE ELEEVEKLVggsHHHHHH | 26.84902727 |
| **36-O51** | MgTQAELERVLAQAKALLAKAEALAEEVLAI LEQIEASPNPTLELLKELGRKTRKLLKAAEAL ASALIAVFRLLGAPQSELDALWQEVLAKLQA VYEATQRAIADKAHLPETRTAVKELLAFLKA QLARAEKLTggsHHHHHH | 86.73654727 |
| **36-O52** | MQAKIEAQLKLAQALVDKAKATAAELRAAV AAIAASPAPTPEQLRAAGELTRKFLKYAEAAT SAITAVYALLGKSKEYQQAIFNETQTLIQAVY EAAQAFLADPSQAPALLAAVDALLATIQKHL DEAKAEAggsHHHHHH | 92.2117426 |
| **36-O53** | MgSQKLLEEQLKQLIEASEKAIKLFDEAIKLA KKLIESPTITEEQKKELGKLVRAAMKALETAY SYLHAVLRLLGKSEEEANAVFNESLEVIKQIY EAAQKALADRSYLPELIELLESRRKELEEYIK EAKKLLggsHHHHHH | 100 |
| **36-O54** | MgREALAKQLLAQAKALLAKAEALAKEVRA LVDKIAADPNAPDSVKRELGVKVRKFVKTAE AHTGAVTAALRLLGVPQETVDKVFNDSLAL ANAVWQAAQEALADRSKLPDFVKAVDDLLK FLQAQLATVEALLggsHHHHHH | 63.22343333 |
| **36-O55** | MgRQALQKELLANAKAALAAVQAAAAAAL ALLDQIAADPAASDELLRALGAAVRAMLKA AETLTGYVVAAYRYIGLPEEEQMAVFNESNA LINAVWEAAQAALADRSAIPAFRQAIADLTA HLTALVARTEALVggsHHHHHH | 109.6155109 |
| **36-O56** | MgTEALLQQLLAQAKAQLAKAQAARDKAL ALLDVIESDPNAPEEVKKQLGKEVRAFLKAA EAATSAITAALRLLGVPQAEVDAVFNENLKLI NEVYEQANEALADRSKVPALKAALDRTLKT LQAQLAKAEKLAggsHHHHHH | 55.31383333 |
| **36-O57** | MEAQKTSLLAQAKAALAKFEEAAAKAKELV | 72.28383333 |

| | | |
|---|---|---|
| | AKIAADPSASDAEKKELGDVVRAMLKAAETALSAITAALRLLGVPEEEVTARFKEGLALIQRVYDAAQAALADPAHAPDFLAAVDELKAFLKAQLAEVEKLAggsHHHHHH | |
| **36-O58** | MEELLKEVLENAEKAVKTAKAKADELLALLAQARADPSAPDSVWRAIGRTFRAFAKAAETAMSYIIAAFRLLGKPQSELDALWQEVTDLIKAAHAALVAAAADRAQLDAAAAQVKALTALLEKYLEEAKKLYggsHHHHHH | 71.69703333 |
| **36-O59** | MEELIKKVIEQAKKLLEEAEKALKELIKLAEEILKSPTVTEEQLKALGEATRKFLKYAEALASAISAAFRLLGKPQEEVQGILDTTLKLIDETYEAAQAALKDRSKLPELIKKAKALLDYNKKQLEEVEKLAggsHHHHHH | 80.03938987 |
| **36-O60** | MgSRALLEELLARADEALATAEAHRDALLALVAEMVAKGEIDPELVKKAGAALRALIRAAETLAGYVIAGFRLLGAPQSELLALHETLTSKIQAVYDAFQAFAKSLTAAPALVAAVDELLAFLRAELARVRALAggsHHHHHH | 41.44313703 |
| **36-O61** | MgREALLERLLEGAEKQLAKFKALAAETRALATALLADPNAPDSVRKALAETTRKMLKYAEATTGYVVAFFREAGKSEEYAQSIWNESLALINAVYDAANAALADRSAIPAFLAAVDKLEAYLKAKVEEAKKLYggsHHHHHH | 80.03938987 |
| **36-O62** | MgREALLARQIAQARALLARAEALAAELKALVERLKAAPEVTDEELRELVRLSRAFLKAFEAALSATLAALRELGAPQEELEALWQAGLELVQAVYDATQKFLADRSHAPELLQAVEDALAYLRAQLARVEEAAggsHHHHHH | 32.25244547 |
| **36-O63** | MgRQALLDSLLEAARAALAKARAQLDVVEALAQAIHDAGTPTPAQLKALGAATRAFIKAAETLNGYILAVYRELGAPEAEIQALHNEFLARVDAVYQAAQAFLADVSQAPAFLAAARDLLAYNEERLAEAEALAggsHHHHHH | 65.263444 |
| **36-O64** | MgRQALLDELLARARELIARAEATAAELLALLARLAADPEPTLEQLLAVRRTLRALLKAAEAAMSAILAAYRLLGVPQEEIDRLWSEHRALIDRVYEAAQRGLADRSHLPEVEAAVAELLAFVRERLREVEELAggsHHHHHH | 37.8934073 |
| **36-O65** | MgTQAQIETTLAAAKAALDAMKAAAEKAAALVDKIAASADVTDEEKRALGAEVRAMLKAAETAISYIGAGLLLLGKPWSEVIALVKETNALVKAVYDAANAFLADRSAAPDFLAAVKALVDFAAEKLKEVEKAIggsHHHHHH | 46.93174943 |

| | | |
|---|---|---|
| **36-O66** | MgTEALLQQLLAQSREFLAKTRALAAEVLALIDRIAASGAPTLEELLELGRLTRALLKTAEALTSSITAALRLLGVPQERVDKVFQEAQKLIQAVYDAANAALATLTALPAFRAAVDALVAYLEAQLKEVEELAggsHHHHHH | 43.57344803 |
| **36-O67** | MgRQAALDRLLALAAAQRAVYDAHAARLVALMDQIYADPNAPDSLYQEAGRVLRAALKAAEAYLSALVAALRELGVPEEEVMAAFNEGLALLNAFAAAAQAFLASRSARPAALAAQAALDAYLAGLEARVRALRggsHHHHHH | 61.07882403 |
| **36-O68** | MgNEEAAKAALAAAEAAAAAVKAAAAEAIALINKIIESGEMKPEDLKALGAAVRAMIKAAEVAVSYVYRLRLLAGVPLEEAVAAAKAALGFITENAEKIREYLKDISKLPNAIELIKAMVAAVGALLAEQKAALggsHHHHHH | 99.99999997 |
| **36-O69** | MgTQATLDAVMAQARALLAKAEALAKELVALAEALAKAKTVTPEQLAALNKALRAFLKAVEAATSAIIAALRLLGASDEEAQAVFDTTRALIQAVYDAFQAFAADRSALPAFLAAVKALLDHVRAQLARAEAAAggsHHHHHH | 70.1843162 |
| **82-O1** | MgEAAALAELRALVAAARARLRELRRAIIKELSDTIEKNKDAPLEEVKKALEEVRDRGVARLEELRAEITAELRALAARVPPSIAVEALAAATELSERVDTESEELVRSTDYILERLEAEKSggsHHHHHH | 53.06103333 |
| **82-O2** | MgAAAELEELRAFARRARAALRDLYRKSILDLSKLLKEVAPLSPEETEARLNALLEKDVAALRALRDAILAELAALAARVSPEVAALAAALAAEITARSDALIADLRRSYDYVLGRILAEKAggsHHHHHH | 14.6299 |
| **82-O3** | MAAELAALRALAREARERIRTTRKALMKELAATAQAAKTLTPAEITAAFDALLAKAKAALDALRTEIRAELDALAATVSPEIQALVAELRAQLDAVIAAEKADFETGIAYVRDQALAAKAggsHHHHHH | 79.28353333 |
| **82-O4** | MgHMEKIIEAGKKLQEERDKVRKVTKENSLAIHDTYTASRPLPAAERIARMNETIAANKEKLTTTYQARIDAAKSVLPLLDEENKKIAEQIIKNTQEAIKQDTAALDTFRARYTAQLQAEAAggsHHHHHH | 57.4545374 |
| **82-O5** | MgAAAELERLRAFVAAARARLRALLRANNLRVTKLAEEVADKTPEEIKAAFDALKAAAAAEFAALLAEIEAEAAALAAAVSPEIAALVREALAQLRADVATEQADFVASVDVLVQRLLDAKAgg | 84.36496667 |

| | | |
|---|---|---|
| | sHHHHHH | |
| **82-O6** | MgSEAELQKIKELLKKLREDFRTLRKQIYADLSALVKEVASATPAEIKAAFDAYRAEATAKFDALAAKAKEELAALAKEVSPEVAKLVKQAEKEITSQITTEQADFNAGVDLVQARRIAAASggsHHHHHH | 5.6875 |
| **82-O7** | MgAEAALAQLRAAARRIRARIAALRKEAMLRLSELTREALTLPVEEVDARFAALRAEVEARFAALEAELTAELDALAKTLPPEYAAQAVALRDEAVARIRAQLADFDRQIDYLRARVVALQEggsHHHHHH | 41.35113333 |
| **82-O8** | MgAEAKLAELRAAIAEDRAAFKELRAAQLKELSAIVTSSASKPLAEYTAALETWLKDFTAKLTALQAEIDARLAALAERLGPEFAPLARAAAAQFKTVIDAEIASLTKGVADLIARKQQVAAggsHHHHHH | 17.827 |
| **82-O9** | MEKELEKLKKLIKKARKEILDLRKEQQKKLSELALANQTKPAEEIEKTLDALKKEAVAKFDELEAKIKKEFEELAKEVSPEVAKLVKEALEELNKLIDTLKKESVDSIDYVRARLLAQAEggsHHHHHH | 84.58133333 |
| **82-O10** | MEEELEKLKALVKEARDYFLTTTRAIQKKLSEVATEAQTLSVEEITAAFDAFRKDAVATLAALRATIKAKFEELAAKVSPEVAKLAKASLAEFNERIATEEKTILDGIDYNKARLVANKEggsHHHHHH | 62.54866667 |
| **82-O11** | MAAELEELRRLIAAARARLRAEVRAAYKALADLSREVLPLPAEEIEKRMDALLAEQTARLDAVKAEDLAALDELAARVSPEAAALAEEARKQLEARWDTAREDLVKGIAYVKARLVAQAKggsHHHHHH | 39.6553 |
| **82-O12** | MgAEAELAAVRELILGAEEEFRAHLKEVQAKMSKELEASLPLPLEEKIKVVKEAYAKAQEEIKKKAEELVARLLAAAAAASPELAALLREAAAQIKERAKAEVEDLKKGEEVLTANYRALLAggsHHHHHH | 42.31626667 |
| **82-O13** | MgAEAAIAELKAKAKELRARITALNKELMKELSALVQQAKTMSAEEVTAAFDQLKTDGNAKYDALAEEQVKELEEFAKAAPEEAKPLAKALAEELKSRIEEQKKEFDKQITYLRDRQVAAKAggsHHHHHH | 48.41813333 |
| **82-O14** | MgSMAELEEVKKLIKEAREKIRELRKEIMLKLSETVKKAKGKSAEEITALFEELKKEVTEKFDELEKEIKEKFEEKAKKVSPELAKLIEEALKQLK | 0.984033333 |

| | | |
|---|---|---|
| | ERIKTEKKDVEKSIDYVRDRLVAQAEggsHHHHHH | |
| **82-O15** | MgSAAELARLRALIDAARAKFREKRKELIKKVTTTATSLLTADPATIEAKFKELKKTVEDELNALITEITAALAALAATLSPEVAALARQAAVEFTNRINTEIKDLNKGLEDLKARLLATQAggsHHHHHH | 71.1116 |
| **82-O16** | MgSAAEEEAIKKAFGEKVRATTKAFEKLMSEAMKELEKLRTKSVEEIKKAKEELKKKILDALKTAVDSIKAAHDEAVKKVSPEDAALIDKITANAVEQLTKLTENFLKQLDARADAIIKEKEggsHHHHHH | 61.23346667 |
| **82-O17** | MgSMAEKEEVEKFIKEKRKEVRAFIKSVYKEISAALTAAIPLPLAEQAKVVEETAAKAIEKLEAKLKEVKKELEEKAAEVSPENAALLLKAGEEFKEIFDGTIADIKKGAAIQVATYTAAAAggsHHHHHH | 63.5077 |
| **82-O18** | MgGEKIVEEAAKLQRELRKELKTLFVAASQKLADTDAAVADAPLAEQVAALKDVAAAEAAKLQAKVDEALKKLEEYYKKVPEEYKPLVKQEIENLKKAGEQDLKALQTLTDRLIATRTAAAAggsHHHHHH | 78.38813333 |
| **82-O19** | MEEELEKLKELAKKGRQKILDTLKEIQKTITKTLKETADKPLEEFEAALNKNLEESLKKLNELLKELKEEIEKLGKTLSPELKELAKELAKDLENIVKVESENLKHSINYLIERRKAEAAggsHHHHHH | 27.49916667 |
| **82-O20** | MgSEEELKLVEELAKESREKFLKLRKKAIKKINEAAEEAAKTLTDEALEKLKATAEEAKKALEEGKKEDEKKAEEIKKKVSPELQKLVEELKKQMKERYDSEKEDIEKSTKYLLERLKKDRAggsHHHHHH | 0.939633333 |
| **82-O21** | MgAAAELAKLKALLDAARAKITSTRKANNTALNTVLTTNASASAAARNAAITAWTATAIANLTALRDEILAEFDALAKTASPETQALLAEAKKEMTARIDAEVTEITASKTDLTARSTAALAggsHHHHHH | 45.7517 |
| **82-O22** | MEEELKKIKELAKELRGKITKLRKAVMKKITETAEEAKTKSVEEIEKLFEELKKEAEKEFDALKTEIKAELDKLKKEVSPEIQKLVTELTKELNSLVDTSLKDINYSIDYVKERLIKEKSggsHHHHHH | 3.253366667 |
| **82-O23** | MgSLAAREELAAALREARANAKAALVAATSELARLELENLTLPLEEGMAALDAAIAAATTTLT | 84.11113333 |

| | | |
|---|---|---|
| | ALLDAERARVEALTERLSPEDRALARAQVETLTKTTRDDLARASTLHRERFRTRATEIAggsHHHHHH | |
| **82-O24** | MgSAAQQARVDELIRQARADLRALLRTLNAAFSEVAFSSESLPLAAYKEALTAAAAAYEATLAAALAAHRARFEALAAQVSPELAARLAEALVQLEAVYTTEAADVRKGVALLIERKEAQAAggsHHHHHH | 83.59886667 |
| **82-O25** | MEAELAELKKLAKELREKIKTLLKKLNLKLYEVATSSRTLSPEEVEAAFAALETTVTTELEALLKESKKKLDALAKKVSPEIAALVDKLKKEIEERINTEKTTLLKGIDYVKARLLAERAggsHHHHHH | 51.61353333 |
| **82-O26** | MgLEAELARLRALVRELREEFRKTLKELNLKISKKAKELSTKDAATIKKEFEKLKEEVRAELEKLLAAHLARLAELAAAVSPEVAALVREAAAEFEHRVETELKALEAGIDYIEARLVAQAEggsHHHHHH | 10.74616667 |
| **82-O27** | MgSAAELAAIKALIDAARARIRALRKEIQRRLTAVLEEIKNLPLAEAKKKLEAYKAEAKAALDALRAEIEAAAAELRAKVSPEAAALVDAALAQLNDVIDAEKAEIDKSIAELLARREKAAAggsHHHHHH | 83.9726 |
| **82-O28** | MgSEAELKLLKEALAAAREKFLALRKQINLDLDKVLKEVAGKSAAEQNAALDKYETDATARLQALAAEITAALQELAAQVSPEYAALFLQAAVEMQARVDAAIADVTTGVADLKARIAAAEAggsHHHHHH | 75.85756667 |
| **82-O29** | MgSEAELAALRELAERARRRFREKLRELHKKLSELLKEVASLPAEEASAAIDKLKEESLKELREFTEELLAELRAAAKTASPEVAALLDELAAELREATETELKALEKSYDSVKAKFLAAKAggsHHHHHH | 100 |
| **82-O30** | MEEKIKEIAEKSRSLLKEIKDIYSKTSQEMADIYLELKDAPLEKIEEALTEKTKEFNEKITAGYDKLIDYLEKAKKEVPPEVQKILDQMIENAKAAKEQELKAINTFKEILLKNARERAAggsHHHHHH | 90.04686667 |
| **82-O31** | MgSAAELAELRARAKAARQKLRDLRKALQLKRSELLQKIQPLSAAEVAAALAKFQAESKAAIEKLLAELTAEFAALAKRVSPEIAALVRALAAELKTIADAEIAEVDTSTAYLTARLLAEKAggsHHHHHH | 4.786833333 |
| **82-O32** | MgDEEQLEKLKKFAKESREKIIKTRKKLLLKE | 1.173033333 |

| | | |
|---|---|---|
| | HEVATEAETASPEEIEKKFEELKKEVEKELNKLKEEIKKKAEEIKKTLNEEYKKLAEEVEKQAKEIIDTEEKTLKEGIDYLKARLLANYAggsHHHHHH | |
| **82-O33** | MEAELAEVRALTRGVRDDFRKLRRAHQLELSALLEAQKPAPAEAKVAALDELTAKFKAEVEAARAADLARIDALQARVSPELAALLAEFRVQVTALYDAELADIEKGAADLKAKIAAEAAggsHHHHHH | 88.66186667 |
| **82-O34** | MEAELEKLRALATQARSDIIALGKENSLLENKVQQQAKTLNAEEAKKAFDDLKTEINAKLDALKADLNAKLDALAAEVSPEVAAKVDVLKTQVNDMINSQKTDFNKGIDVLLANYLALKSggsHHHHHH | 48.15243333 |
| **82-O35** | MgKEKEIEELKKLAKEARKRFIELRKENYLELSKAVKEAQTKSPEEIKKILEKWKEDSTKKLDDLLKKIEAELKKKAAEVSPEIAKLVKELAAQMKVVFDAEKEDIEKGVKEVIKQLTSAASggsHHHHHH | 1.080966667 |
| **82-O36** | MAAELAELKELIKSSREKFIATRKENQLAFNKVLTELADKPAEVRVAALEKVAAQAKENLAKLLKEIEKELKELAAKSSPEIQKLVEEALKQFKEIVDTEQKNVDTQLKDLVARIKAVDAggsHHHHHH | 18.90273333 |
| **82-O37** | MgSEAELERLRALARRVREELSALVKEINLKIHEVSTKVRSASPEEIKKAFDELLKEAKAKLEALRERQLAELEAAARTASPEARALLRELGAQATERVDAEIEDITKGINEIVERLTKVASggsHHHHHH | 69.89226667 |
| **82-O38** | MgVAAELAALRALLKAARDAFTALLKAINAKLSATVQAVLNADAETIRAALEAVRDEALAAIDALTAAQLAELDALAAAVSPEVAALVREAAAQIAARGATERADFEKSVDEILARLVSRAEggsHHHHHH | 101.6064667 |
| **82-O39** | MgRAAELAELEALAKEIREDFVKTIKEQNAKLSKLLQETKNADLATKTAAIKAWEKEAKAALDKLLAEHEARLAELRARVSPEVAALVDELAEQARERVDAQKEDFAKGAKEIIARITAEAAggsHHHHHH | 77.23926667 |
| **82-O40** | MgSEAELAKVKELAKEFRDLVLKLIKEQTLKEYEAFSAAETLSAAEAKATFDALKTELTAAFNAVKDEIAAKAAELKAKLSPELAALVAEIEKEANERVDTNLKDFLKGIDIRLATYVAAKSggsHHHHHH | 88.29973333 |

| | | |
|---|---|---|
| **82-O41** | MgLEAALARLRALAAEFREALKKKRKEAVLLLTKTATESATLSAEEAEKKFAALREKVTAIFDALKAEVEARAAALRAEVPPEVQPLVDELLAQALERVTTEKESLLKGIDYVKARILAQAEggsHHHHHH | 1.3251 |
| **82-O42** | MgAEAEQKQLEELAKNSREDYVNLRKEENARFAAVVEAVRGKSPEEIRAALTAELEKITDRLKKLKADNEKKITTAAAKVSPEIAAQLKKLAEQVKDYVDSAIKEAKKSTEIVTAKLEAAAAggsHHHHHH | 64.85303333 |
| **82-O43** | MgVEEELEKLKKLVREAVANFRALLKAINAKLTEVVTAVLGKPASEIVSALDAYLDAALAELNALTEAEVERLREFRKTVSPEVAKLADEAIEELRARAETEAADLKKSVAYLKERLLATSAggsHHHHHH | 88.3069 |
| **82-O44** | MgGAEDLALLRAAARESLAAVRALRKALVLKLTEAAKAAETKSAAEVTASFAKLKAEVKAEFDALAAELAARLAALAARVGPAARPLVAALAAELQTRIDTEKAALLKSIDYLLARLLAVKAggsHHHHHH | 2.7367 |
| **82-O45** | MgSSAAEAARRALAAETRKRIHQIVKDLNAKRSKAIEEVKSLPAEEIRKKLEELKKEDVAELKAGLDAIRAEAVARLRAIPGVTPEEIAAMEAQLAAIRDTLTKDIEKSYDDLIERLTAEAEggsHHHHHH | 91.3628 |
| **82-O46** | MgSEAQLAEVRALIAATREDIRTLRRALIAQFSALATAALTLPAAEVTKKFAAFKATVTAAFDARAAAVTTRFDEVAARVSPEIAELVLAARDELLERITTERADLLTGIAEVEKRLTAHNAggsHHHHHH | 91.09153333 |
| **82-O47** | MgAEAELARLRELARGTLAELRALRRELQRRLTEVVEKSKTLSAEEVQKLFDAFKEEATAAFDALRARVTARLEAAREELSPEFRPLVEELAREMDARIDTERADFLRSTDELLKRALAEKEggsHHHHHH | 76.52726667 |
| **82-O48** | MgTEAVRREVAAFLREARAKVKKLVVKASKDMADIYKKNKDKPYEVVKKELDKVLNEALKELDKILKETEEFVAKKAPLLSPEDQALLAQEVANMRAAKEQDTKALQTHRDNLLALAKKESEggsHHHHHH | 99.38753333 |
| **82-O49** | MgSAAELAKVKAFAKKARAEIRATFKELNKKLAEIATEGQTLPPEEIEAKLNALKTDATKTVTAKLNALKAEGEALAKTLSPEQKALVDAIITELTARANTELADFKKSVDYVKERLLAAQAggsH | 100.1883667 |

| | | |
|---|---|---|
| | HHHHH | |
| **82-O50** | MgRAADLEKIRALVYAQRQALLTLRKELNRKLHEVLTSNASAPVEKQVAALEEYLKDATARLQALRAASRAQLEALAATLDPEAAALVREAIAQITAQIDAEIADFEKGVAELKKRIRAAAAggsHHHHHH | 37.7989 |
| **82-O51** | MAAALEALRALVRAARDRLRTVIKEIQLDISKTVEEVRDKPAEEVRAALTSLRDEGVARIDALVAAVEAELAAFAATAPPEIQALAAEAIAQVRAVADAEKADITTGIDEILARLLAEKEggsHHHHHH | 83.9427 |
| **82-O52** | MEEQLARLRELARAGREALRKARKAAYAAISATLTATAGASLAEFEAALTATVAAAKAALDKVLAENRAALEALAAEVSPEIAALARQAAAELAKVAAALKEDLDAGEKELIARRRAVAAggsHHHHHH | 100.0000333 |
| **82-O53** | MgAEAELAALRELAAEVRAAITQTKKDSSLKIYETSTANKGKSVAEFKAALTALAEQIAAEIDAKLAEVNARLDALAATVSPEIQALIAELRAQATTAFTTLAADTKAGIADIIARYAAIAEggsHHHHHH | 88.5669 |
| **82-O54** | MgSEAELAELRALAKEIREAFVKLVKDYSLRLNKVLTENKDAPVEERVAAIEKLRDDYKARLDALAAEYTARLEELAARASPEVAALARALAAEVRDRAATLKADVDKTTAVILERVKKEASggsHHHHHH | 81.85626667 |
| **82-O55** | MEEELEKLKKLVKEARERFRDLRKELNLELNKVLTAVQTKSPEEITATLAKELERIEKRLEEERKKIREEFEKLKKEVSPEVAKLVEEAIKQLEERFRVEAERYRKGLAILTARLVANRAggsHHHHHH | 38.6861 |
| **82-O56** | MgAEAALARIRELARAARADITTLLRKSNLRLSELLKEVKDLPLEEAEAKLKKLETDDKAELDALAAALEARAAALRAELPPEFQVLVDALAAELRARITTEKAALSKQYADILARIKANRSggsHHHHHH | 54.90796667 |
| **82-O57** | MgSAAALAAIRARVAAARQRVVDLRRELVLALSKAAEAVKDLPAEEVRKVFDALRVELAARFDALAAEIEAEFDALEAEVPPEFQALVREARAQILEVIATEKEDLLAGVDYVLARLVAEKEggsHHHHHH | 82.22526667 |
| **82-O58** | MgTEEAKKQAGAILKDTQKTLKATIVAADNEIATYLKENAAASLAEKKAHVDEILKATAAEVDKILADAKAKLDALLPKLTPEDQAIVTSQLAT | 65.49396667 |

| | | |
|---|---|---|
| | IESTAKEELEKRKTYWKQKVEAEDAFNKggsHHHHHH | |
| **82-O59** | MgAEAERAELRALGLAGRAALTTAFKQHQKALHETAEAVRDASAAEIRAALDALRAQAKADIDTLLAAQLAELEAFAARASPEVAALARELAAELKAIADAMKADIDKSVDVVLSRLLARAEggsHHHHHH | 76.27426667 |
| **82-O60** | MEAELEKLRKLAKEGRKRIIDLRKKHNLKLSEKIKEVKTKSAEEIKEELEKLKEEQKKEFEKLKEEIFAELDELAKTASPEIKALLDELKKDLEKYYNTELEDLDKSTEYLTERLVKEAEggsHHHHHH | 0.836333333 |
| **82-O61** | MgSEAALERIRALIAEADARIRDLLKELNLEVSRVATAAQTLSAAEIDAAFAALVERVGARLDALAAEIEARLAALAATLPPEHRKLVEEAIRQLTGRIATERADFGTRLAVLRQRLLAQNAggsHHHHHH | 63.03033333 |
| **82-O62** | MgSEEALKKIKELAKEARERIVALQKEASLALSKTAKAVADKSAAEVTAALDALTTEVTAKFDALKAELTKELEALKKEVPKEYQPLVDELKKQILAMVDAQKADLTKGIADVKARMVAVKEggsHHHHHH | 11.61206667 |
| **82-O63** | MEKELEVLKKKIKELRKKVIELYKELLKEYNKILKEVLNKSAEEIKKALEEAEKKFKEKLEALAKEIEAELEALAKKFSPEIAKLAKEAAKQLKERIESMIKDFEKGSKILLERLTAQAEggsHHHHHH | 6.903933333 |
| **82-O64** | MgSAAWEAELRALARASREAITALLKEANRKLSELAKEVYNKPAEEIKARFEALKKEFLAKFDALEAEIEARIAALRERAPPEFRPLLDELAAQSRERIAVEKEDLVKSIDYVLEKVLAVAEggsHHHHHH | 66.96093333 |
| **82-O65** | MEEQLALLRQAARAARARARELFREANLLVHEAATAALTLPPEEVRALFAALREEVTAKFEALKAELTAELDALAAQVSPEVAALYEQLKVEVSGVIDTELADFDKGIAYLEARRLALLAggsHHHHHH | 39.67263333 |
| **82-O66** | MEADLERLRALLREARERFKALRKEIWGELAKAVQEAKTLSAEEAAALFDKVLAESLARLEALRAEDEAALAALRAAVSPEVAPLVDEAAEQLNKYYDSFTEELRRSIELVRQRVLDAKAggsHHHHHH | 0.7641 |
| **82-O67** | MgSAAELARLRAATAEALARIRATLKELVKKLSEVAKEAQTKSIEEAKKLFEEVKEYARERLS | 74.99323333 |

| | | |
|---|---|---|
| | ALRTEIEAELAALAATLSPEYQALAAAAAAE LSGRIDTELERLKYDANYVLERYEKEKKggsH HHHHH | |
| **82-O68** | MgREAELAELRRLVAESRAAIRALRKEAVLM LTKAIEESKTLSPEEVKKLFEDLKTEIAARFD ALAAEIEARFRALAARVSPEIAALALEAAAQ MRDVLATEKADLLKGIDYVLKRILAEKEggsH HHHHH | 3.212933333 |
| **82-O69** | MgREAELAELRALAAEIRKARRALLKEIQLET NKIIKEVMNATPEEIKAKLAALEKEQLARVQ ALRDDFLARVRALAARASPEIAALADQLAEQ IETDFAIDEADIKKSIADITAKLLAKAAggsHH HHHH | 66.3891 |
| **82-O70** | MgSEAALERLRAAARELRQELLDTIKAEWLK LTELVTANASAPLETRVAALQAQQAAILAKL DAKGKEIAARLARLAEELGAEFAPLVEQLGV ELQKIIDETKKDIEKGVAELIARAKAEAAggsH HHHHH | 104.6864 |
| **82-O71** | MgSQAELEKIKEFAKQVREDLVALYKEHQLR LTKIATEKASAPPAEITAALDALEKEFKAELQ ALLDKQLAKLDEMEKTVSPEHQPLVTELKK QAQEIADTLLADITKSVDDVRAKLLAAYAggs HHHHHH | 96.9003 |
| **82-O72** | MgAAAATAAVRALILGSRKELRELLKKLQLK FSELVRSVIGLPAEEIKEKLEKLLEEIKKELEK KVKELVARLAAAAETASPEDAALLAEAAVQL TSEAETELKDFEKQFEYVTSRLIAQAKggsHH HHHH | 100 |
| **82-O73** | MEADLARLRELARRIRERELALIKRLNKKISE LTKEVAGKPAAEVEKAFDALAKEAEAEFAAL LAEIEAELAELEREVSPEVRPLVRELRAQARE RIETNLAAIKKGIDYLKARKLKEAAggsHHHH HH | 0.366433333 |
| **82-O74** | MgDEAYLAAVKELIATNRARITALRKAVLLDL TALTEKVKDADAATVKAALDALAAAAQARF DALAADIEAEAAALAAQVPPEYAALVKAAL EQLKARIDALKAETAKTIQTVRDRMLAEKTg gsHHHHHH | 3.544866667 |
| **82-O75** | MgSMAEIEELKKLAKEVRDEIVALLRAGQRD LTAVLKETANADAATRVAAVRAVRADRLARL DALEAKILARLEELKKSLSPEAAKLAEKLGE EVKAIVETEKADIDRSTAVLEERLKADASggs HHHHHH | 79.17123333 |
| **82-O76** | MgTQALLEAHAAEIRAGLKARKAAFTAAMS | 51.40446667 |

| | | |
|---|---|---|
| | AMAAAAAAAATKSPADIDATLAAATAAAKAALDTTTQAVLDALKKAAEELPAEYQPLVKQMGENVKAARTTETKAIDTFAARLRANALAVAAggsHHHHHH | |
| **82-O77** | MgLEAALAALRALVAEVRAKFTATRKAIYKALSDAVKAVQTQSAEEAKATLETVKADGVAALKDLRATTTAAILEFAATLPEELRPLALELAKQFTEYTDTEIKDLTKGYDEVLARVLAQKAggsHHHHHH | 0.296766667 |
| **82-O78** | MEAELELLRKKAKEVRQRFLELRKENNLKVNEVAKAVLNKSPEEIKAELEKVRDEIKADLEALLAEIKAELEKLAKEVSPEIAALVKELAAEMEVYVKSEAEAFEKGIAELTKRLVAQAEggsHHHHHH | 0.807366667 |
| **82-O79** | MEAEVERLRALARAARAELVTTLKAINNKLSKTAAAVRNASPAEATAAFNAVKTDATAALTALAARHEARLAALAAEVSPEVAALARELAAELRARIDTEKTNIGKGIDYVLKRVLAEVAggsHHHHHH | 0.4914 |
| **82-O80** | MgREEELALLRERIRAARERLRALRREAQRKLAELVEEVKSLSPEEITKAFDELKKEVEERFDALLAAVEEEFRALAATLSPEIAALADEAIAQLREVIETERERLLKSIEEVKERVLAAKAggsHHHHHH | 19.8024 |
| **82-O81** | MgAAAATAELAALQKAALDAYRKALKAFQKEVSTTLEETKPLPAEERLKILTEVIEKGEEDLLKIIDQGIADLKAALEKHRSLLSPETIATVNAQIEGFERAKEEVKKSSAVLKEKIKATVEggsHHHHHH | 85.92573333 |

| **Screening mini-protein sequences for mIL-4Rα** | | |
|---|---|---|
| **Binder No.** | **Sequence** | **BLI-Response** |
| **1** | MSEAEEKKKAAQKKLEEAAKKARELTGKGRKLLKDPSTKEEGEKLIDKGLQIEADAAEEYIEVTKAVEKAggsHHHHHH | 0.0083 |
| **2** | MSENRELYLEIRGKLQALETTAYAGTEEEAKEAVKKAVELAKEYGSEEYVKATKENMEKLAKIGIERRKggsHHHHHH | 0.0783 |
| **3** | MNLEEAQKKALEKAEEAVALAKEYGLYKPEYLEEYLKKIKEIAAKSLQSAEAALAEAELFKERVEELYKKIggsHHHHHH | 0.0117 |
| **4** | MEKRKKFLNDRIEYTSEYLKELADKKPNAEELKKFIDEGKEKALEANEKDPELGEETLQ | -0.0095 |

| | | |
|---|---|---|
| | EMNAEAMLKINggsHHHHHH | |
| **5** | MEEKLKEARKRYEEAREEIRKKEAETLEKNSSLSPEEIKAKVLEVSEKLTKEAVEKIEKEVKEggsHHHHHH | 0.0052 |
| **6** | MDLLKDAEAAAKELKKELEEAKELAKENRKKAAYLAYSVEQQAKDLADYYKSEGDKESAKLMEEVSKEAKKVYEEIAKggsHHHHHH | 0.008 |
| **7** | MPDIEKAAKEAKERVLESFEKLKDNPELLKLDVQMLIERYRELDPRLEEAAKRGALEAAKEVAKDPAAAAIKEAALEggsHHHHHH | -0.0237 |
| **8** | MSVEALIEEAKKQMEKEKEIAKKLAEKDKEYGLQSVKQAEDSLTYRAKETGKRVGLSEEEIEKVIKVIKEKAKEYKKEIEKggsHHHHHH | -0.0023 |
| **9** | MDKEKLEAMAKAAEKYGEVAGKEMAKIDPSLAEAAKYQGIASAYYRIAKIASPEDAKELLAKAKEADEKAAALIEggsHHHHHH | 0.0588 |
| **10** | MLEKYLERTKEAVKEMAKISPEYALLEGEMAKERMKLLAEKVSPEAAEKLKKLGEEVEKYAKELAKEKAggsHHHHHH | -0.0094 |
| **11** | MSLEEKAKKVAEEVLAALEEGKKKALEAYKTEGLEAAKKVWDEAREKAQTLLTELVEESPEAAKIAVKIIQARSDAKAEAggsHHHHHH | 0.026 |
| **12** | MSEKVEEQKKAVDEVAKEIAEAQKEVHEKAKNKEEAAKLLKEKSEEFEKKGKEEKNEYASAAYLQAADHFNYRAKKVKEggsHHHHHH | -0.0207 |
| **13** | MEEEERRKKLEEARKKLSELQGKVIEGAEKSLEEAKKYIKELAEEAKKYGSEEYAKELEKESLKAAEELHKLNKggsHHHHHH | -0.0034 |
| **14** | MNEELEKGKKEIAKAGEEGAKELVAAPKEKFKEVLAKVARRLADLANEAAKKIGSKEAAKELREARYEAEGAAFGAGIDALggsHHHHHH | 0.0101 |
| **15** | MKEEAVKKYVEKVAEDTKPLPLEYQELNLETLAARFELIDPELGKVAEETGKKELEKIKKERggsHHHHHH | -0.0072 |
| **16** | MEEKLKKAKEVVDKISKEGREEAIKEAKKVAEEKGKEAGLKKYEEMRWETQKKEADAYAEIIEggsHHHHHH | -0.0025 |
| **17** | MSGEKLLESVKKRMEEEYKIAKELAKKDKEYGLQQIKQAEYSLKYRAEEVGKKLGLKEEDIEKVLKEIEKLAKEYEEEIKKggsHHHHH | 0.195 |

| | | |
|---|---|---|
| | H | |
| **18** | MSKEVIEKGLEYIREARNYALGLKKKNPEKYEEIMDKILEAERKLFENPEEGLKLAKEVKEEAKKLLEggsHHHHHH | -0.018 |
| **19** | MSEKLEEGKKEIKELGKEYAKKLVAASPEDFKKVLDEASKALRDKANEKGKEIGNKEEAKELREVAYEEEGKAFNAGIDAKggsHHHHHH | -0.0158 |
| **20** | MSNEELAKEAAEKALELRKKGIEEAEKAYKTEGLEAGKEVYAKYRDEIQKTISDLLEKNPEAAKIAVEILQAESDKEAKEggsHHHHHH | -0.0057 |
| **21** | MKEKAEKSLKLGEEYYKKGLEYEKEGNYKEAAYYYSKAARYYAKAMELYEKVGDKEKAEEAYKKYKEAMDKTYELLKKINggsHHHHHH | 0.0001 |
| **22** | MVEKLKEEMEKAEKEALEVGKEAREAYLLAEKAKKEDPEKAKKYEEEADKLYQKERELTLKAYELKKEYEKAKKELLEggsHHHHHH | 0.0093 |
| **23** | MEKKKIIAYYLYRGYRLVTGLNAIGAKYGQPEIAKEASVKVWEKYIIAAKAAIKNKEEGEKLAEEADKIAADAAYKIYQAAggsHHHHHH | 0.3215 |
| **24** | MSERREELERAREAAREAEAKVYNYMIDNPSATKEEALELAKKLAEEAGKKLKNPEAAKDAVKMALEGSKIVIEKYFAYKggsHHHHHH | 0.0017 |
| **25** | MKEEEAEKLLLAADAYTAAGYVIEDPNDELAKLKMEHYLKKLEGEAKEEVLKAIEEAKKLSKEEGQEYLTDLAIEYIKKAKggsHHHHHH | 0.4575 |
| **26** | MKEKTERFMKEAEKRAEEAAKIMEELGGDPIQAKAQAYDSFAYRAEKLGFTEAAKVFKKKSEEYTAKALAggsHHHHHH | 0.0276 |
| **27** | MEEEKIKKAKEEVKKISEKARKEVNEALYLAKKDPSTAEKWLAKADEAQKEESDKILEYEKIKKggsHHHHHH | -0.0011 |
| **28** | MSEIEELLKKAAEYYAKAVEDPSNRAKYIAIGDAYMRKARELELKEEREKRLKggsHHHHHH | -0.0179 |
| **29** | MKEEIYKETREEMLKISEEMRKKLSEEAKKAPPEKRKEVIEKGREEIRKKETELAEEANKKVEggsHHHHHH | -0.0182 |
| **30** | MEEEKKKLLEEAELYKEAGEAEYKMALERGEIEAKKLEEKDPEKGKKALEQYEKVGEE | -0.0173 |

| | | |
|---|---|---|
| | LKKKYEEKAKELEEKAKKggsHHHHHH | |
| **31** | MKEKIEKAKEFREEVRDKANAAAAEAAEKVSPEEAKEIALKEGEKAIEESNKYYRSLLggsHHHHHH | -0.0175 |
| **32** | MSLEELKKKEEEEKRKKAKEKRDEIYDEAGKAAADAITADPANAATKAKAIQDKGFEAARAYEKELKEggsHHHHHH | -0.008 |
| **33** | MDLLKDAKEAAKELKEEVKKAKELAKEDPKKAAKLAYRVMEQAKDLAKYYESLGDKESAELMKKVSKEAEEAYKEIKEggsHHHHHH | 0 |
| **34** | MSRREELRKKRDEIYDEGNAKAAKAVEEDPENAKEKAAAISQEAFERARELEREggsHHHHHH | -0.005 |
| **35** | MEEEEKRKKYLALAKYYRKGAEEYRKLAKEAAADPKLKQSAEGAENTAKYLEEEAEKLEKEAKggsHHHHHH | 0.0187 |
| **36** | MAEIEKKKELLAKAAEYERKAYSGNKEEAIENGEKAAEYYKEVGYEEYAEYVLNRVKEYAKKKggsHHHHHH | 0.0303 |
| **37** | MSEKDKGRIALGDEYGAKGKALLAAGGDLNEAAYYFGRAARYYNAVPSTQAKADEAYAQYADLTLAAAKAAggsHHHHHH | 0.0248 |
| **38** | MEELEKEMAKRLEQGKKAKEAMEKIHKKTYEIGKKHKDKKEALEESAKLAEKAAEENKDNEYAYANMQMAADRLKELAKggsHHHHHH | -0.0051 |
| **39** | MDAKEIYVETAKETAKEIKALAEKEKDPEAKAKIEELAKKAEKAAKEAAKSTEEEAKKNYEIAGNALREAEAIAKKggsHHHHHH | -0.008 |
| **40** | MAMEHYLAEAKKLDAKGTLKDKALAYDNYAGAYRRLAEYYEKLGKKEEAAKYNALAEKYNAKSQEYATAYRDSggsHHHHHH | 0.1171 |
| **41** | MSSTEEEIKKYAEELAKKAEEAVGPDPSAKRHARESLRITAENALAAGKSVEEAKELLEAEAQMVVERYEILKKRAAggsHHHHHH | -0.0102 |
| **42** | MSEALKKALEEAKKESAEKIKETDKIAKEYMKKHSPEEGKKFAKELLLKTRDEIQKETRNKLEEVRggsHHHHHH | -0.0075 |
| **43** | MTKEYAKRTLETAKELAEKAIKEGRYDVAEAVYLQAADGFERRAKEAKDEEAKKFLLEQAEKAKKLAEEVKKKAAggsHHHHHH | -0.0112 |
| **44** | MSEENKKLAEYYEAAAEAEKKLGEELLKEAKEKDLPEKERKLLIERAKRAFEDAKVFEE | 0.0315 |

| | | |
|---|---|---|
| | KAKELKAKggsHHHHHH | |
| **45** | MSALEEERKKLLKKGKYYEYRAKRAKEDGEKLSAELKAKGDEELAKNVKEAYDLYVKEAEAEAKKYKAAAEAggsHHHHHH | -0.0017 |
| **46** | MDERVKKYYEETYKKAKEAVEILKKNPKSTVEMLEAEAIQAGYGYADRAKTEEEKKAALEAAKEVAKEAKKEWAggsHHHHHH | 0.0144 |
| **47** | MSKEELEKVFKTNSEKMREAKYEYLRVKKTGTPEEAKEAEKKRDEILTELNDKIVDAYEEYKKSggsHHHHHH | -0.006 |
| **48** | MKAEAIRKYAEREAEAVRALAKEAAAKSPELAKTLAEGSLAQGRARAEELAKIDPEGGKIYREIVEATAAELLAAggsHHHHHH | -0.0076 |
| **49** | MLNEKEKLEIIGEASYSLAKELLEKAKKGEANISDLASGYRAAEEAAKKFEELGNEEKAKEAKKIAEELKKLAKggsHHHHHH | 0.0036 |
| **50** | MEEKKKLIEKAEELYDKALEIEAKEGNAEAMYNLSKAYGLYRKGKDLETAEKLAEKASPEAKKYLEEAAEYFKKAKEggsHHHHHH | 0.0061 |
| **51** | MEELEKEAEKKLGEAHDLFYEAGTSGNLTPEVQEKIRELGNEARRLRLLYEKKKKEEggsHHHHHH | 0.0229 |
| **52** | MALEKEKEELKKKALELYDKAGRAYREAEIKGNKEAAELALKSMEYATEAYYSIDNIEKSKEYLEKAEKYLKEAEKLLEggsHHHHHH | 0.037 |
| **53** | MDLELKAEGYEREKKALKAAKELEGEEGVENYYKWKAESVMERAKIAGIKVTFEEALELAKKREEELEEAYKKLKEggsHHHHHH | -0.0008 |
| **54** | MSENEETRKKILEYSEKAVSEANKAAKEEGMEAAIKVAEEKAKEAEKLDEYIGRDAKMILERIKYLASggsHHHHHH | 0.0045 |
| **55** | MTEEYAKRTLEEAKKLAKKAKAAGRWDVAEAVYLNAAAGFEERAEKTPDPEHKKYLLEKAKEMKELAEKVKKEAAggsHHHHHH | -0.0015 |
| **56** | MEWREKARAARDEIRDKVGAASAAAIDADPAKAKETAAKLAEEGRAAANEVARALLAEAEAAggsHHHHHH | 0.01 |
| **57** | MEEIEELLKKAAEYYEKAYEDPENRAEYIKLGDYYARKARELELKEKVLKKLEggsHHHHHH | 0.0017 |
| **58** | MDPAEIIAQAQAEMKEAAKKYLKKAQEEGKSLSPEERKKLLEEYRIKAQEEQRKIAEKAREKIAggsHHHHHH | 0.0127 |
| **59** | MLNNKEKKEVLLEQAKIAAEESKKYKELA | 0.0084 |

| | | |
|---|---|---|
| | EKYKSNPELYKKYALEAQRAERGAQAYQAIADELAAKIggsHHHHHH | |
| **60** | MNEAQKALEEAKKEAEKAKEAAEKAKKEIGLKKLSERAIYKINSLVNFAESLASNPEKALIAAEAAKEYAKEEKEKFEKLLggsHHHHHH | -0.0006 |

| **Alanine mutants used for crystal screening** | | |
|---|---|---|
| **Binder No.** | **Sequence** | **Binder No.** |
| **82-O32-native** | MgDEEQLEKLKKFAKESREKIIKTRKKLLLKEHEVATEAETASPEEIEKKFEELKKEVEKELNKLKEEIKKKAEEIKKTLNEEYKKLAEEVEKQAKEIIDTEEKTLKEGIDYLKARLLANYAHHHHHH | 82-O32-native |
| **82-O32-M1** | MgDEEQLEKLKKFAKESREKIIKTRKaLLLKEHEVATEAETASPEEIEKKFEELKKEVEKELNaLKEEIKKKAEaIKKTLNEEYKKLAEEVEKQAKEIIDTEEKTLKEGIDYLKARLLANYAHHHHHH | 82-O32-M1 |
| **82-O32-M2** | MgDEEQLEKLKKFAKESREKIIKTRKKLLLKEHEVATEAETASPEEIEKKFEELKKEVEKELNKLKEEIaKKAEEIKKTLNEEYKKLAaEVEaQAKEIIDTEEKTLKEGIDYLKARLLANYAHHHHHH | 82-O32-M2 |
| **82-O32-M3** | MgDEEQLEKLKKFAKESREKIIaTRKKLLLKEHaVATEAETASPEEIEKKFEELKKEVEaELNKLKEEIKKKAEEIKKTLNEEYKKLAEEVEKQAKEIIDTEEKTLKEGIDYLKARLLANYAHHHHHH | 82-O32-12 |
| **82-O32-M4** | MgDEEQLEKLKKFAKESREKIIKTRKKLLLKEHEVATEAETASPEEIEKKFEELKKEVEaELNKLKaEIKaKAEEIKaTLNEEYKKLAEEVEKQAKEIIDTEEKTLKEGIDYLKARLLANYAHHHHHH | 82-O32-13 |
| **82-O32-M5** | MgDEEQLEKLKKFAKESREKIIKTRKKLLLKEHEVATEAETASPEaIEKKFEELKaEVEKELNKLKEEIKaKAEEIKaTLNEEYKKLAEEVEKQAKEIIDTEEKTLKEGIDYLKARLLANYAHHHHHH | 82-O32-14 |

| **Arginine mutants and fusion proteins** | | |
|---|---|---|
| **Binder No.** | **Sequence** | **Binder No.** |

| | | |
|---|---|---|
| **82-O32-A12R** | MgDEEQLEKLKKFrKESREKIIKTRKKLLLKEHEVATEAETASPEEIEKKFEELKKEVEKELNKLKEEIKKKAEEIKKTLNEEYKKLAEEVEKQAKEIIDTEEKTLKEGIDYLKARLLANYAggsHHHHHH | 82-O32-A12R |
| **82-O32-L111R** | MgDEEQLEKLKKFAKESREKIIKTRKKLLLKEHEVATEAETASPEEIEKKFEELKKEVEKELNKLKEEIKKKAEEIKKTLNEEYKKLAEEVEKQAKEIIDTEEKTLKEGIDYrKARLLANYAggsHHHHHH | 82-O32-L111R |
| **82-O32-E100R** | MgDEEQLEKLKKFAKESREKIIKTRKKLLLKEHEVATEAETASPEEIEKKFEELKKEVEKELNKLKEEIKKKAEEIKKTLNEEYKKLAEEVEKQAKEIIDTrEKTLKEGIDYLKARLLANYAggsHHHHHH | 82-O32-E100R |
| **82-O32-43c** | MgDEEQLEKLKKFAKESREKIIKTRKKLLLKEHEVATEAETASPcEIEKKFEELKKEVEKELNKLKEEIKKKAEEIKKTLNEEYKKLAEEVEKQAKEIIDTEEKTLKEGIDYLKARLLANYHHHHHH | 82-O32-43c |
| **82-O32-47c** | MgDEEQLEKLKKFAKESREKIIKTRKKLLLKEHEVATEAETASPEEIEcKFEELKKEVEKELNKLKEEIKKKAEEIKKTLNEEYKKLAEEVEKQAKEIIDTEEKTLKEGIDYLKARLLANYAHHHHHH | 82-O32-47c |
| **82-O32-58c** | MgDEEQLEKLKKFAKESREKIIKTRKKLLLKEHEVATEAETASPEEIEKKFEELKKEVEcELNKLKEEIKKKAEEIKKTLNEEYKKLAEEVEKQAKEIIDTEEKTLKEGIDYLKARLLANYAHHHHHH | 82-O32-58c |
| **82-O32-72c** | MgDEEQLEKLKKFAKESREKIIKTRKKLLLKEHEVATEAETASPEEIEKKFEELKKEVEKELNKLKEEIKKKAcEIKKTLNEEYKKLAEEVEKQAKEIIDTEEKTLKEGIDYLKARLLANYAHHHHHH | 82-O32-72c |
| **ABD035-82-O32** | MgLAEAKVLANRELDKYGVSDFYKRLINKAKTVEGVEALKLHILAALPggsggsggDEEQLEKLKKFAKESREKIIKTRKKLLLKEHEVATEAETASPEEIEKKFEELKKEVEKELNKLKEEIKKKAEEIKKTLNEEYKKLAEEVEKQAKEIIDTEEKTLKEGIDYLKARLLANYAHHHHHH | ABD035-82-O32 |
| **ABD035-82-O32-43c** | MgLAEAKVLANRELDKYGVSDFYKRLINKAKTVEGVEALKLHILAALPggsggsggDEEQLEKLKKFAKESREKIIKTRKKLLLKEHEVATE | ABD035-82-O32-43c |

| | | |
|---|---|---|
| | AETASPcEIEKKFEELKKEVEKELNKLKEEIKKKAEEIKKTLNEEYKKLAEEVEKQAKEIIDTEEKTLKEGIDYLKARLLANYAHHHHHH | |
| **82-O32-A12R** | MgDEEQLEKLKKFrKESREKIIKTRKKLLLKEHEVATEAETASPEEIEKKFEELKKEVEKELNKLKEEIKKKAEEIKKTLNEEYKKLAEEVEKQAKEIIDTEEKTLKEGIDYLKARLLANYAggsHHHHHH | 82-O32-A12R |
| **82-O32-L111R** | MgDEEQLEKLKKFAKESREKIIKTRKKLLLKEHEVATEAETASPEEIEKKFEELKKEVEKELNKLKEEIKKKAEEIKKTLNEEYKKLAEEVEKQAKEIIDTEEKTLKEGIDYrKARLLANYAggsHHHHHH | 82-O32-L111R |
